# *Anopheles stephensi* as an emerging malaria vector in the Horn of Africa with high susceptibility to Ethiopian *Plasmodium vivax* and *Plasmodium falciparum* isolates

**DOI:** 10.1101/2020.02.22.961284

**Authors:** Temesgen Ashine, Hiwot Teka, Endashaw Esayas, Louisa A. Messenger, Wakweya Chali, Lisette Meerstein-Kessel, Thomas Walker, Sinknesh Wolde Behaksra, Kjerstin Lanke, Roel Heutink, Claire L. Jeffries, Daniel Abebe Mekonnen, Elifaged Hailemeskel, Surafel K Tebeje, Temesgen Tafesse, Abrham Gashaw, Tizita Tsegaye, Tadele Emiru, Kigozi Simon, Eyuel Asemahegn Bogale, Gedeon Yohannes, Soriya Kedir, Girma Shumie, Senya Asfer Sabir, Peter Mumba, Dereje Dengela, Jan H Kolaczinski, Anne Wilson, Thomas S Churcher, Sheleme Chibsa, Matthew Murphy, Meshesha Balkew, Seth Irish, Chris Drakeley, Endalamaw Gadisa, Teun Bousema, Fitsum G Tadesse

## Abstract

*Anopheles stephensi*, an efficient Asian malaria vector, recently spread into the Horn of Africa and may increase malaria receptivity in African urban areas. We assessed occurrence, genetic complexity, blood meal source and infection status of *An. stephensi* in Awash Sebat Kilo town, Ethiopia. We used membrane feeding assays to assess competence of local *An. stephensi* to *P. vivax* and *P. falciparum* isolates from clinical patients. 75.3% of the examined waterbodies were infested with *An. stephensi* developmental stages that were genetically closely related to isolates from Djibouti and Pakistan. Both *P. vivax* and *P. falciparum* were detected in wild-caught adult *An. stephensi*. Local *An. stephensi* was more receptive to *P. vivax* compared to a colony of *An. arabiensis*. We conclude that *An. stephensi* is an established vector in this part of Ethiopia, highly permissive for local *P. vivax* and *P. falciparum* isolates and presents an important new challenge for malaria control.

**Summary of the article:** *An. stephensi*, a metropolitan malaria vector that recently expanded to the Horn of African, was highly susceptible to local *P. falciparum* and *P. vivax* isolates from Ethiopia and may increase malariogenic potential of rapidly expanding urban settings in Africa.

## Background

With expanded global malaria control efforts there have been two decades of substantial declines in malaria cases and deaths. These successes were mainly attributable to wide-scale deployment of vector control tools and availability of efficacious treatment [1]. Control programs in Africa traditionally focus on rural settings, which is where most infections occur [2] although malaria transmission is also a health concern in some urban settings [3, 4]. In 2015, 38% of Africans were living in urban settings; the number of Africans residing in urban areas is expected to double in the coming 25 years [5]. Urban settings can be sinks of malaria transmission primarily associated with importation of malaria from (rural) areas of intense transmission due to movement of people at the urban-rural interface [6]. With the adaptation of existing vectors to urban environments [7] and emerging vectors such as *Anopheles stephensi* in urban areas [8], malaria transmission in urban settings is becoming more likely. Urban areas can thereby form foci of active malaria transmission [9]. *An. stephensi* is an efficient vector for both *Plasmodium vivax* and *P. falciparum* in Asia and is the dominant malaria vector in India and the Persian Gulf [10]. *An. stephensi* predominantly breeds in urban settings with a preference for human-made water storage containers [11]. Recent reports indicate that *An. stephensi* is spreading in the Horn of Africa (Djibouti [13], Ethiopia [14] and the Republic of Sudan [15]). *An. stephensi* emergence has been epidemiologically linked to an unusual resurgence in local malaria cases in Djibouti city [16]. A recent technical consultative meeting convened at the World Health Organization (WHO) identified that there is potential for spread of *An. stephensi* across Africa and urged for more data on its distribution to allow monitoring of potential spread of *An. stephensi* from the currently affected areas and on the vector’s susceptibility to local *Plasmodium* isolates [15]. In the present study, we examined the abundance of *An. stephensi* in an urban setting in Ethiopia, characterized its aquatic habitats, biting and resting behavior, and, for the first time, examined its competence to transmit local *P. vivax* and *P. falciparum* isolates.

## Methods

### Description of study site

This study was conducted in Awash Sebat Kilo town (916 meters above sea level; 8°58’59.99” N 40°10’0.01” E), on the main transportation corridor from Addis Ababa (220km southeast) to Djibouti (Figure 1). The town has an estimated total population of 24,700 [17]; the semi-arid climate is dominated by a major rainy season (July-August) and short intermittent rains (April/May) and an average temperature of 25.8°C (17.3°C-33.6°C) [18]. The Awash River Valley is the most irrigated area in the country with extensive river-fed agriculture. Malaria transmission is perennial in the area surrounding the town with annual parasite incidence of 536 per 1000 population in 2019 (five-year trend summarized in supplemental notes). Entomological surveys conducted in 2018 detected the occurrence of *An. stephensi* in Awash Sebat Kilo town [19].

**Figure 1.**
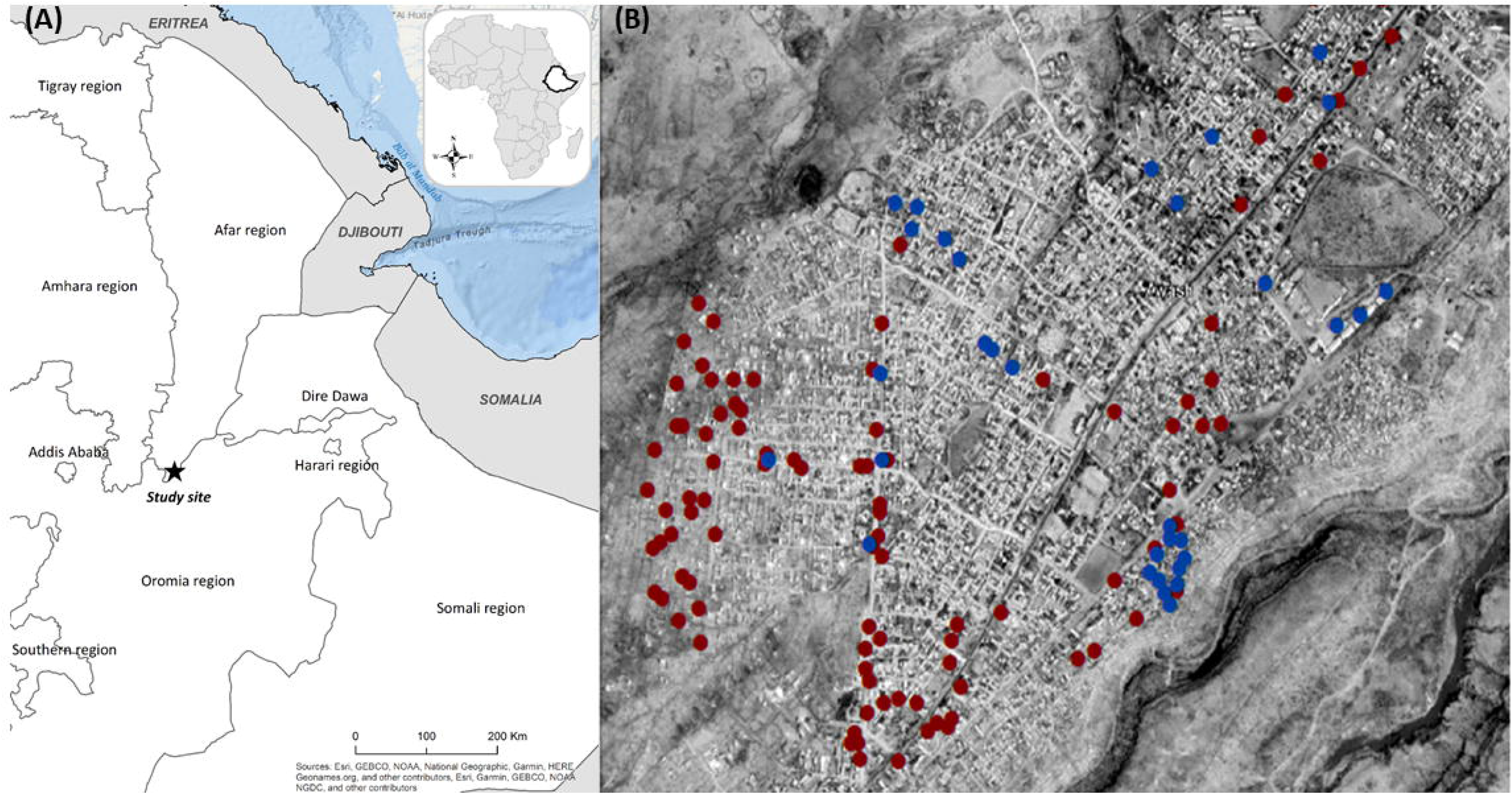
Map of study site, *An. stephensi* aquatic habitats and adult mosquito resting sites. Indicated are the map of Ethiopia with regional boundaries and study site (starred) (A) and an aerial view of Awash town (B) with larvae/pupae (red dots) and adult mosquito collection sites (blue dots).

### Characterization of aquatic habitat, resting, feeding and biting behavior

Aquatic sites were examined for the presence of *Anopheles* developmental stages by standard dipping (10x) for 5 consecutive days. Developmental stages were separated from culicines in the field and transported to Adama malaria center for rearing to adults. The resting, feeding and host-seeking behavior of *An. stephensi* was assessed using five conventional entomological sampling techniques: i) Centers for Disease Control (CDC) miniature light traps (Johns W. Hocks Company model 512) catches, ii) human landing catches (HLC), iii) pyrethrum spray sheet collection (PSC), iv) aspiration from animal shelters, and v) cattle-baited traps (Supplemental notes). Adult mosquitoes were identified morphologically using standard identification keys [20]. Fully fed mosquitoes identified as *An. stephensi* were kept in paper cups at Adama laboratory in ambient conditions and allowed to lay eggs on filter papers soaked in water on a cotton roll for egg ridge counts (Supplemental notes).

### Mosquito rearing and membrane feeding assay

*An. stephensi* were reared from larvae/pupae (from aquatic site examinations) to adult at ambient temperature (26±3°C) and relative humidity (70±10%) and fed with fish food (Cichlid Sticks; King British Fish Food, Tetra). *An. arabiensis*, the principal malaria vector of Ethiopia, from an established colony were reared under identical conditions and maintained with 10% sucrose solution. Following informed consent, patients who presented to the Adama malaria clinic with microscopy confirmed *P. vivax* and *P. falciparum* mono- and mixed-species infections were asked to donate venous blood sample (5mL) in lithium Heparin tubes (BD Vacutainer^®^). Asexual parasite and gametocyte densities were quantified by two expert microscopists on thick blood films prepared from finger prick blood samples, screening against 1000 leukocytes. Thin blood films were examined to identify *Plasmodium* species.

Four-to-seven day old adult *An. stephensi* and *An. arabiensis* were starved for 3 (*An. stephensi*) or 12 hours (*An. arabiensis*) before feeding. One hundred and twenty mosquitoes of each species, 40 in each of 3 paper cups, were fed fresh patient blood through membrane in the dark for exactly 25 minutes (Supplemental notes). Fully fed mosquitoes were maintained under the same laboratory conditions with 10% sucrose solution for 7 days post feeding before being dissected for oocyst detection and for 12 days for sporozoite detection.

### Molecular detection of parasites and blood meal sources and targeted sequencing of morphologically identified *An. stephensi* mosquitoes

*Plasmodium* infection status of individual wild-caught morphologically-confirmed adult *An. stephensi* mosquitoes was assessed using nested polymerase chain reaction (nPCR) targeting the small 18S subunit [21] using genomic DNA extracted from homogenate of mosquito’s head-thorax and abdomen separately [22], indicating sporozoite and oocyst-stage infections, respectively. Multiplex PCR that targets the mitochondrial cytochrome b gene and produces species-specific fragments of varying sizes was used to assess blood meal sources of individual mosquitoes [23]. For confirmation of morphologically identified *An. stephensi*, DNA was extracted from whole mosquito bodies using the DNeasy Blood and Tissue kit (Qiagen, UK). PCR was performed for each individual mosquito, targeting the nuclear internal transcribed spacer 2 region (ITS2) and the mitochondrial cytochrome oxidase subunit 1 gene (COI) [24]. Following PCR clean-up (Source BioScience Plc, Nottingham, UK), chain termination sequencing was performed to generate unambiguous consensus sequences for each sample (Supplemental notes). Sequences were assembled manually in BioEdit v7.2.5 [25] to create unambiguous consensus sequences for each sample. Consensus sequence alignments per gene were generated in ClustalW and used to perform nucleotide BLAST (NCBI) database queries [26]. *An. stephensi* ITS2 and COI sequences, from across the vector’s geographic range, were downloaded from GenBank for phylogenetic analysis in MEGA X [27]. Additional outgroup ITS2 sequences were retrieved for *An. maculatus, An. maculipalpis, An. sawadwongporni* and *An. willmori*. Alternate maximum-likelihood (ML) phylogenies were constructed using the Jukes-Cantor (ITS2; final tree *ln*L=-916.913) or Tamura-Nei (COI; final tree *ln*L=-732.248) models, following appropriate nucleotide substitution model selection in MEGA X. Bootstrap support for clade topologies was estimated following the generation of 1000 pseudoreplicate datasets.

### Statistical analysis

Analyses were performed in STATA version 13 (StataCorp., TX, USA) and GraphPad Prism 5.3 (GraphPad Software Inc., CA, USA). Feeding efficiency (proportion of fully fed mosquitoes) was compared in matched experiments using the Wilcoxon matched-pairs signed-ranks test. Logistic regression was performed to compare infection status between *An. arabiensis* and *An. stephensi* using individual mosquito data and a fixed effect for each human participant to account for correlations between mosquito observations from the same donor. Bland-Altmann plots were generated for differences in infectivity between mosquito sources with Pitman’s test of difference in variance.

## Results

### Most of the potential aquatic habitats were infested with *An. stephensi* developmental stages

Eighty-five water bodies within Awash Sebat Kilo town were assessed for *An. stephensi* larvae and pupae. All of these water reservoirs were human-made (Figure 2; Supplemental notes). *An. stephensi* larvae were detected in 75.3% (64/85) of sites (Table 1; Supplemental notes); of which the final aquatic developmental stage (pupae) were detected in 37.5% (24/64) of the waterbodies. Larvae were more commonly found in permanent (85.4%, 41/48) compared to temporary containers (63.9%, 23/36; *P*=0.022). The most common water body co-inhabitants were *Aedes aegypti* (39.1%, 25/64) and culicine mosquitoes (23.4%, 15/64). A total of 49,393 immature *Anopheles* larvae and pupae were collected in 20 visits for rearing; of which 45,316 (91.7%) emerged to adults. Morphological identification of 1,672 female *Anopheles* confirmed that all were *An. stephensi*.

**Figure 2.**
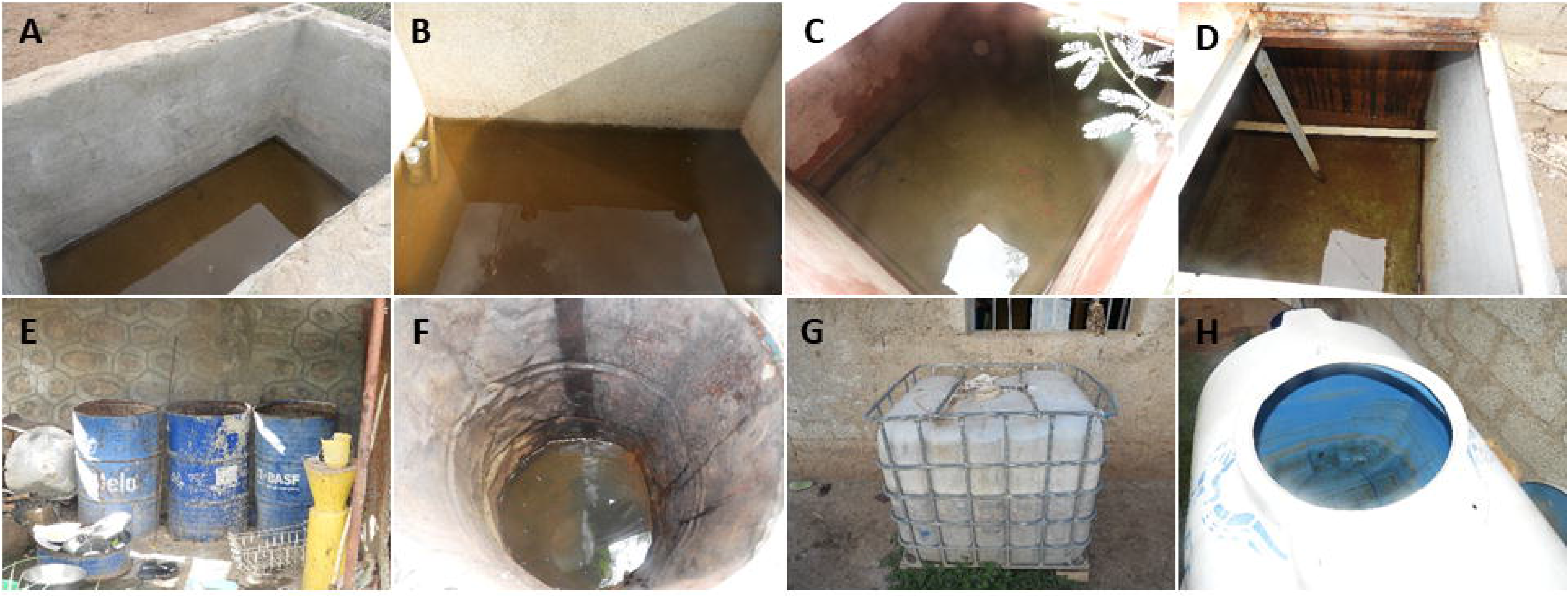
*An. stephensi* larval habitats. Images are of waterbodies that were infested with developmental stages of *An. stephensi*, namely water reservoirs made of bricks or cemented tanks (A – B), custom-made metal containers (C and D), barrels (E – F) or plastic containers (G – H). The median volume of the aquatic containers was 4m^3^ (IQR, 1.0 – 15.6) and ranged from 0.06m^3^ to 360m^3^. The majority of the containers were uncovered and were in use for household (32) and construction purposes (34). The material from which the different types of reservoirs were made of included cement (n=45), plastic (n=9), fiber (n=14) and steel (n=17).

**Table 1.** Characteristics of the aquatic habitats surveyed

| **Characteristics** |  | **Habitat**<br>**s** | **Larvae**<br>**detected** | **Pupae**<br>**detected** |
| --- | --- | --- | --- | --- |
| **Localities**<br>**within the**<br>**town**<br>**(Kebeles)** | Sebat Killo | 60 | 73.3% (44/60) | 43.2% (19/44) |
|  | Lemlefan | 17 | 70.6% (12/17) | 0% (0/12) |
|  | Alalamo | 8 | 100.0% (8/8) | 62.5% (5/8) |
| **Water body**<br>**type** | Permanent | 48 | 85.4% (41/48) | 41.5% (17/41) |
|  | Temporary | 37 | 62.2% (23/37) | 30.4% (7/23) |
| **Shade status** | Fully shaded | 22 | 63.6% (14/22) | 42.9% (6/14) |
|  | Partially shaded | 24 | 99.7% (22/24) | 31.8% (7/22) |
|  | Not shaded | 39 | 71.8% (28/39) | 39.3% (11/28) |
| **Usage** | in use | 71 | 76.1% (54/71) | 37.0% (20/54) |
|  | not in use | 14 | 71.4% (10/14) | 40.0% (4/10) |
| **Container**<br>**material** | Fiber jar/tyre | 23 | 43.5% (10/23) | 40.0% (4/10) |
|  | Metal (steel<br>tanks/drum/barrel) | 17 | 94.1% (16/17) | 31.3% (5/16) |
|  | Cemented/Ceramic | 45 | 84.4% (38/45) | 39.5% (15/38) |
| **Cleanliness** | Clean water | 56 | 80.4% (45/56) | 37.8% (17/45) |
|  | Turbid water | 28 | 67.9% (19/28) | 36.8% (7/19) |

### Adult mosquitoes rest mainly in animal shelters and feed also on humans and are infected with *Plasmodium*

A total of 89 adult female *Anopheles* mosquitoes, the majority of which were blood fed (72), were collected in two monthly rounds (6 days each) of entomological surveillance (August and September 2019) with a median of 10 *Anopheles* mosquitoes per productive trapping night (range 1-22). The majority (80.9%, 72/89) were morphologically identified as *An. stephensi;* the remainder were *An. gambiae* (n=16) and *An. pharoensis* (n=1). Most of the *An. stephensi* mosquitoes were collected from animal shelters (91.7%, 66/72); the remainder (8.3%, 6/72) were collected outdoors using HLC (Supplemental note). Almost half (43.8%, 7/16) of the *An. gambiae* were caught by CDC light traps, but no *An. stephensi* mosquitoes were caught by this method. Of the adult caught *An. stephensi*, for two non-blood-fed mosquitoes the abdomen was positive for *P. vivax* (2.2%, 2/89) indicating oocyst level infection and one blood-fed mosquito collected from an animal shelter was positive for *P. falciparum* (1.1%, 1/89). From blood meal analysis the majority of adult *An. stephensi* had fed on animals (52.8%, 38/72); such as goat (n=23), cow (n=7) and dog (n=5) with a non-negligible number of them feeding on humans (12.5%, 9/72) (Supplemental notes). A quarter of them (23.4%, 11/47) fed on multiple sources including humans.

### *An. stephensi* are highly susceptible to infection with Ethiopian *Plasmodium* isolates

A total of 47 paired membrane feeding experiments were conducted using blood from patients with microscopy confirmed *P. vivax* (n=36), *P. falciparum* (n=7) and mixed *P. vivax* and *P. falciparum* (n=4) infections (Table 2). The majority of patients were female (73.8%, 31/42) with a median age of 27 years (IQR, 19-38). Gametocytes were detected by microscopy in the majority of *P. vivax* mono-species infected patients (73.5%, 25/34) but fewer in patients with *P. falciparum* (14.3%, 1/7) and mixed species infections (25.0%, 1/4; only *P. vivax* gametocytes). A total of 4,088 female *An. stephensi* raised from field collected larvae/pupae were fed alongside age-matched 6,130 colony derived *An. arabiensis*. The proportion of blood fed mosquitoes was generally higher for *An. arabiensis* (median, 80.5%; IQR, 72.5-85.0) compared to *An. stephensi* (median, 53.5%; IQR, 44.0-68.0; *P<0*.001; Figure 3A).

**Table 2.**
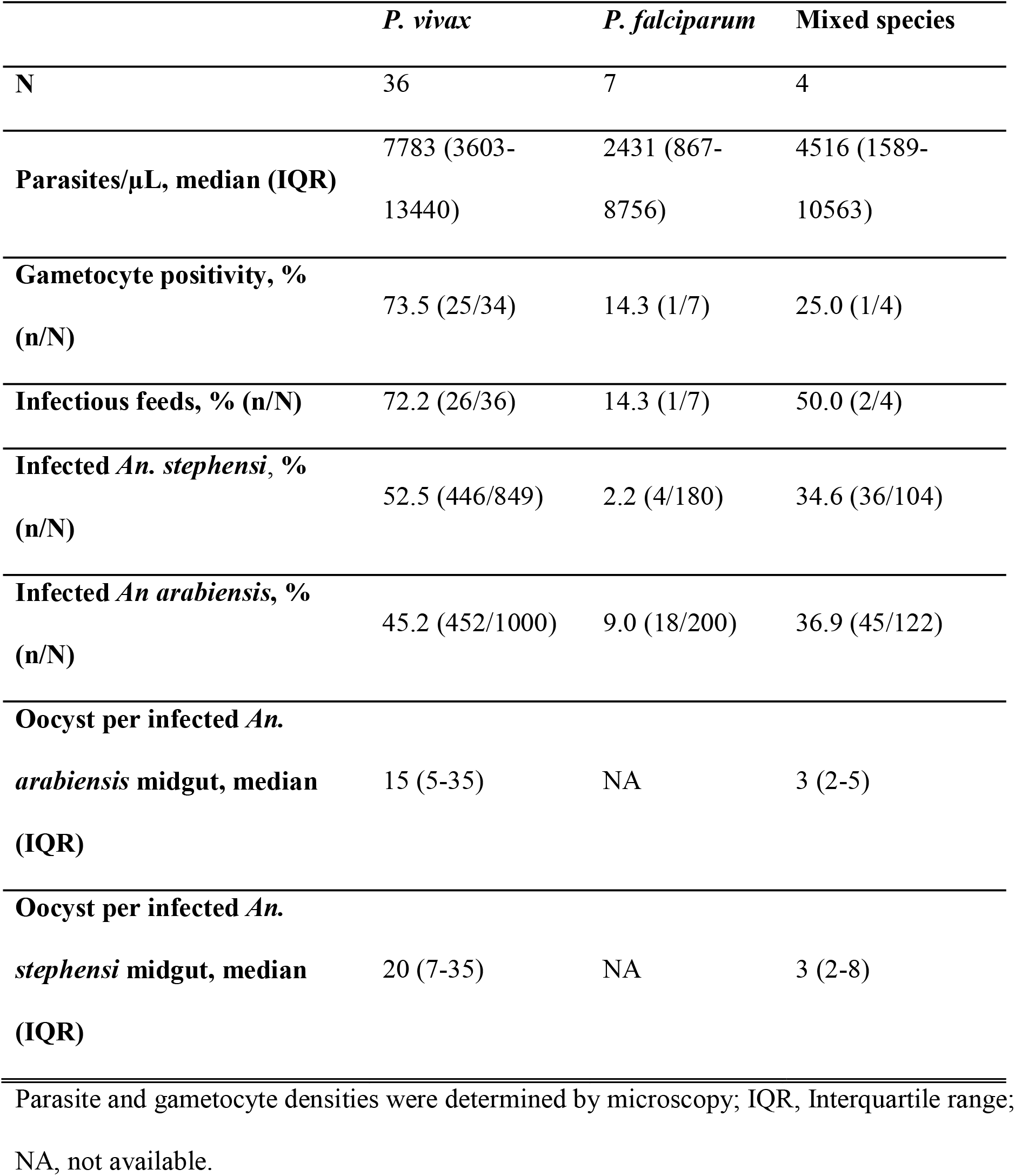
Membrane feeding assays: characteristics of malaria patients and mosquito feeding outcomes.

**Figure 3.**
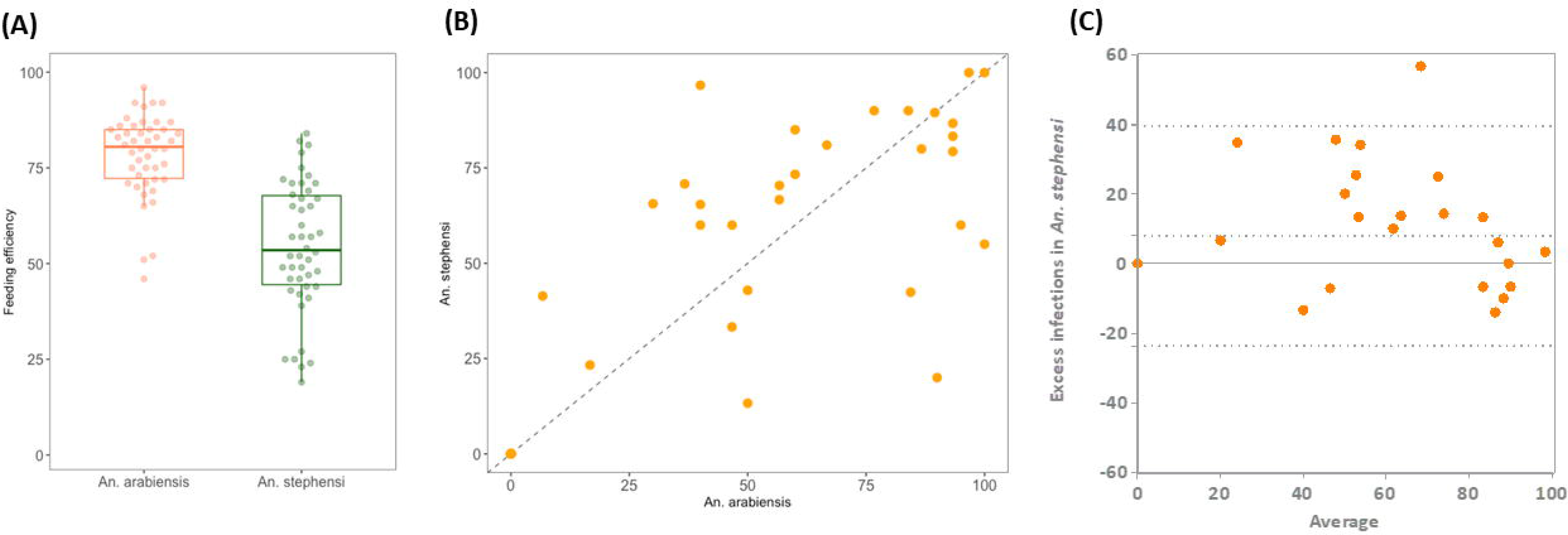
Comparison of feeding efficiency and infection rates for *An. stephensi* and *An. arabiensis* in paired feeding experiments. The percentage of fully fed mosquitoes for *An. arabiensis* (red circles) and *An. stephensi* (green circles) (A). The percentage infected mosquitoes for the two mosquito sources (An. *stephensi* on the Y-axis and *An. arabiensis* on the X-axis) (B). The Bland-Altman plot (difference plots) for mosquito infection rates in different mosquito species (C). Symbols indicate the difference in infection rate in *An. stephensi* versus *An. arabiensis* (Y-axis) in relation to average infection rate in these two species (X-axis). Positive values (57.1%; 16/28) indicate a higher infection rate in *An. stephensi;* dotted lines indicate the 95% limits of agreement. There was no evidence that correlation coefficient between the paired differences and means differed significantly from zero (Pitman’s Test of difference in variance, r=0.026, *P*=0.864).

For each blood feeding experiment, an average of 24 (range, 10-33) *An. stephensi* and 28 (range, 19-32) *An. arabiensis* mosquitoes were dissected for oocysts on day 7 post feeding. Overall, 72.2% (26/36) *P. vivax*, 14.3% (1/7) *P. falciparum* and 50.0% (2/4) mixed species infected patients infected at least one *An. arabiensis* and one *An. stephensi* mosquito. A very strong association was observed between the proportions of the two mosquito species infected with *P. vivax* (ρ=0.82, *P<0*.001; Figure 3B) with a statistically significant higher proportion of infected mosquitoes for *An. stephensi* (median, 75.1%; IQR, 60.0-85.9) compared to *An. arabiensis* (median, 58.4%; IQR, 40.0-85.6; *P<0*.042). Allowing for the number of dissected mosquitoes for each set of paired feeding experiments results in higher odds of infectivity to an individual mosquito for *An. stephensi* (Odds Ratio [OR], 1.99; 95% CI, 1.52-2.59; *P<0*.001) (Figure 3C).

Oocyst intensity per infected midgut was higher for *An. stephensi* (median, 17; IQR, 6-33) than *An. arabiensis* (median, 13; IQR, 4-30; *P<0*.001; Figure 4A). Oocyst intensity associated positively with oocyst prevalence for both *An. stephensi* (ρ=0.553, *P<0*.001) and *An. arabiensis* (ρ=0.576, *P<0*.001) mosquitoes (Figure 4B). To further determine competence for transmission, random subsets of blood-fed mosquitoes from six paired feeds were kept until day 12 post feeding for sporozoite quantitation in salivary glands. Sporozoites were detected in both mosquito species and higher sporozoite loads were detected for mosquitoes from batches where oocyst prevalence and intensity were higher (Supplemental notes). Among paired feedings, after accounting for number of examined salivary glands, the odds of detection of sporozoites was substantially higher in *An. stephensi* (OR, 4.6; 95% CI, 2.2-9.9; *P<0*.001) compared to *An. arabiensis*.

**Figure 4.**
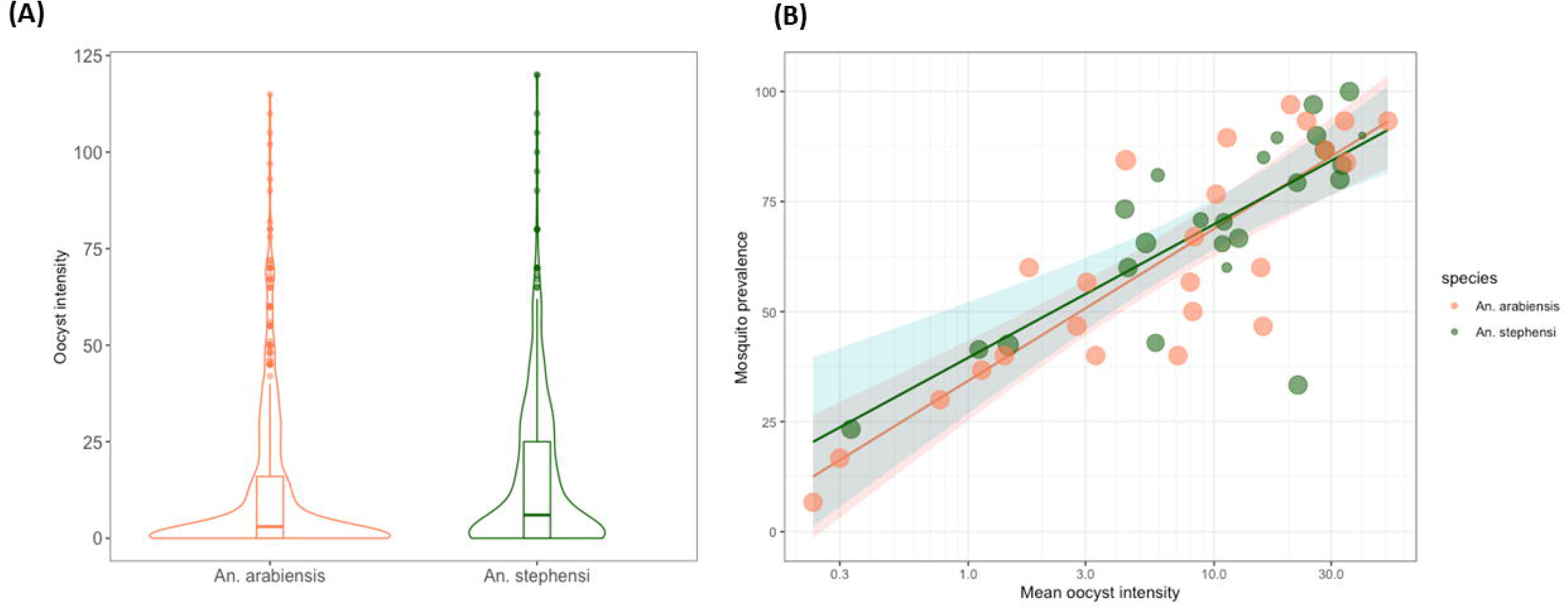
Comparison of oocyst intensity and prevalence for *An. stephensi* and *An. arabiensis* in paired feeding experiments. Oocyst intensity (number of oocysts per dissected midgut) for individual mosquitoes of each of the two species (A). The violin plot presents the estimated kernel density, the median is indicated with horizontal lines, the interquartile range by the box and upper and lower-adjacent values by the spikes. In panel B, oocyst prevalence (proportion of midguts with detectable oocyst) (Y-axis) is indicated in association with Log_10_ transformed oocyst intensity (X-axis) for *An. stephensi* (green dots) and *An. arabiensis* (orange dots). Data are presented for 24 feeding experiments where 723 *An. arabiensis* and 643 *An. stephensi* were dissected.

### Sequencing confirms Ethiopian *An. stephensi* mosquitoes are closely related to *An. stephensi* from Djibouti and Pakistan

DNA extracted from 99 mosquitoes representing all larval habitats was used for determination ITS2 and COI sequences. Of these, 76 were successfully amplified and sequenced for ITS2 while 45 were successfully amplified and sequenced for COI. All of sequences were confirmed to be *An. stephensi*. The ITS2 phylogeny was constructed from a consensus alignment of 301bp, containing 124 variable sites; the COI phylogeny was constructed from a consensus alignment of 465bp, containing 17 variable sites. The ITS2 tree indicated that *An. stephensi* from Ethiopia form a well-supported monophyletic clade (bootstrap value of 100%) with all other *An. stephensi* sequences from across the Arabian Peninsula and South-East Asia (Figure 5). The COI tree was more resolutive, suggesting *An. stephensi* from Ethiopia were most closely related to mosquitoes from Djibouti (64%) and Pakistan (54%). Four haplotypes using COI and two genotypes using ITS2 were detected.

**Figure 5.**
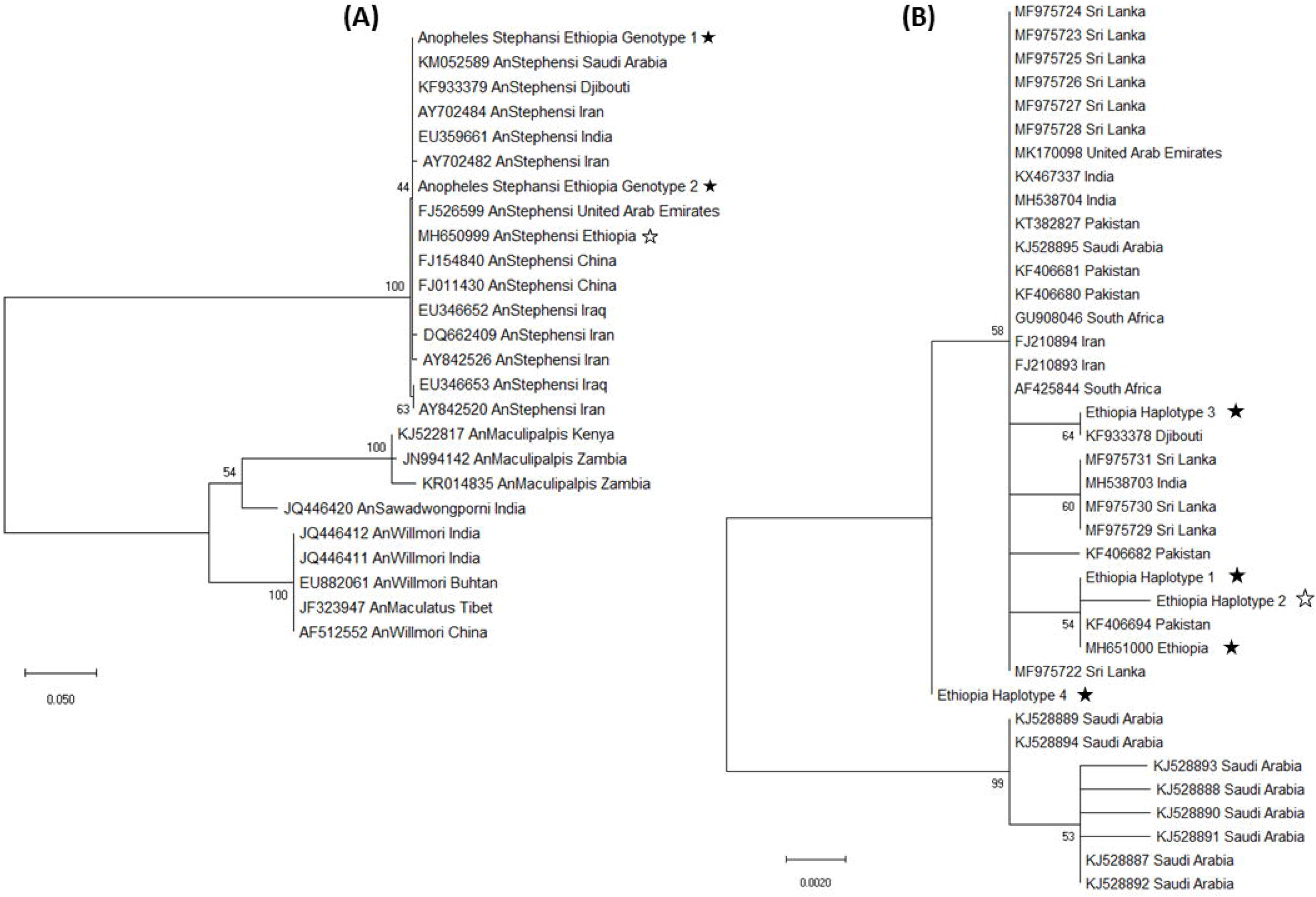
Maximum-likelihood phylogenies of ITS2 (left) and COI (right). Maximum-likelihood topologies were constructed using representative reference sequences with published geographical data downloaded from GenBank. Within the Ethiopian population, due to the presence of a hyper-variable microsatellite region, ITS2 sequences (A) were trimmed to create a consensus alignment of 289bp; one polymorphic site separated samples into two genotypes (indicated with filled asterisk together with the previously reported genotype, MH650999, Carter, et al. [14] in unfilled asterisk). COI sequences (B) were assembled into a consensus alignment of 687bp; a total of four variable sites were identified, corresponding to four haplotypes (indicated with filled asterisk together with the previously reported genotype, MH651000, Carter, et al. [14], unfilled asterisk). Nucleotide sequences for ITS2 and COI were deposited in GenBank under the following accession numbers: Ethiopia Genotype1, MN826065; Ethiopia Genotype2, MN826066; Ethiopia Haplotype1, MN826067; Ethiopia Haplotype2, MN826068; Ethiopia Haplotype3, MN826069; and Ethiopia Haplotype4, MN826070.

## Discussion

In this study, we examined the abundance, behavior and vector competence of *An. stephensi* in an Ethiopian town, Awash Sebat Kilo. *An. stephensi* was the dominant vector, larvae being present in the majority of examined human-made water bodies. The detection of *Plasmodium* developmental stages in adult *An. stephensi* demonstrates its receptivity to local parasites. This was further demonstrated by mosquito feeding assays where *An. stephensi* more frequently became infected and infectious when feeding on blood of *P. vivax* patients compared to an insectary adapted colony of *An. arabiensis*. These data demonstrate the widescale presence of a novel efficient vector in this urban area in Ethiopia.

Originally reported in India, *An. stephensi* has expanded westward from the Persian Gulf, documented in farms and within the capital city of Kuwait in 1981 [28] and subsequently in the Riyadh region of Saudi Arabia in 2007 [29]. More recently, it spread into the Horn of Africa where it was reported in Djibouti city in 2013 [13] and Ethiopia in 2016 [14]. The recent emergence in the Republic of Sudan [15] and more widespread sites in Ethiopia [19] in 2019 suggests the species has the potential to become a widespread African malaria vector. Our data demonstrate that *An. stephensi* is firmly established in an urban setting in Ethiopia located on the main transportation corridor from Djibouti to Addis Ababa. The detection of four haplotypes using COI and two genotypes using ITS2 suggests the independent arrival of different populations or heterogeneity arising after the importation of the mosquito species. Our findings further corroborate recent suggestions that *An. stephensi* in Ethiopia is closely related to populations from Pakistan [14]. Regardless of its origin, it is evident from our data that the mosquito is abundantly present; of the 85 water bodies examined, 64 were infested with developmental stages of *An. stephensi* even in the driest months of the year (May/June), further indicating how well-suited the mosquito is to local weather conditions and the availability of human-made water storage containers. The number of larvae/pupae we detected (~50,000 in twenty rounds of sampling) and the development to adulthood (>90%) is an alarming confirmation of adaptation in this setting.

Uniquely, we directly determined the vector competence of wild-caught *An. stephensi* to naturally circulating *Plasmodium* parasites from malaria patients via membrane feeding in comparison to an established and membrane-adapted colony of *An. arabiensis* [31]. Our mosquito feeding experiments predominantly included *P. vivax* clinical cases who are highly infective [31, 32], and allow a sensitive comparison of mosquito species. Although the membrane adapted colony of *An. arabiensis* had high feeding rates, mosquito infection rates were statistically significantly higher for *An. stephensi* than for *An. arabiensis*. Our detection of salivary gland sporozoites establishes that sporogonic development of local *P. vivax* can be completed by *An. stephensi*. We recruited fewer clinical *P. falciparum* cases who, in line with other findings, were less likely to infect mosquitoes compared to *P. vivax* patients [31]. Despite a modest number of observations, our findings demonstrate that also local *P. falciparum* isolates are capable of infecting *An. stephensi*. This *ex vivo* evidence of susceptibility to local *Plasmodium* isolates is further supported by the detection of adult mosquitoes infected with *P. falciparum* and *P. vivax*. This is, to our knowledge, the first direct evidence of infected *An. stephensi* in Ethiopia.

The spread of *An. stephensi* can be linked with movement of goods and people [13] and the favorable conditions created by rapid social development and urbanization [33] that is accompanied by increased availability of aquatic habitats in the form of water storage tanks. Rapidly expanding urbanization often leads to informal settlements with poor housing and sanitation [34]. Although housing conditions are improving in Africa, particularly in urban settings, there are still major gaps such as poor estimates of combined water, sanitation and hygiene coverage [35]. Establishment and potential spread of *An. stephensi* in the Horn of Africa poses considerable health risks of increased receptivity and local transmission of malaria in the increasing urban African settings requiring realignment of malaria control programs. Its frequent presence in human-made aquatic habitats [37] indicates that a simple bifurcation between urban and rural settings may be misleading and is context dependent. Additionally, urban populations are also at increased risk of *Aedes*-borne diseases, which are increasing in incidence in Africa [38–41]. We regularly detected developmental stages of both *An. stephensi* and *Aedes* in the same water body [42]. Outbreaks of chikungunya [43], dengue [44], and yellow fever [45] were recently reported from Ethiopia within the same geographic setting. The WHO recommends the use of integrated vector management [46] – an adaptive and evidence-based approach to vector control which utilizes vector control interventions from within and outside the health sector that may include larval source management of both *An. stephensi* and *Aedes* vectors. Our findings support that larval source management may be considered to prevent further spread of *An. stephensi* and Aedes-borne disease outbreaks in African towns and cities and beyond their territories [47]. Further investigation is required to understand how *An. stephensi* respond to the existing and novel insecticides and vector control strategies.

## Declarations

### Ethics approval and consent to participate

The study protocol was reviewed and approved by the Institutional Ethical Review Board of the Aklilu Lemma Institute of Pathobiology of Addis Ababa University (Ref.No. ALIPB IRB/025/2011/2019), the Oromia Regional Health Bureau (Ref. No BEFO/MBTFH/1331), and AHRI/ALERT Ethics Review Committee (Ref.No.AF-10-015.1, PO07/19). All participants provided written informed consent; parent/legal guardians for participants younger than 18 years. Those collecting human landing collections also provided written informed consent and were monitored for 3 weeks following collections and treated if any malaria symptoms occurred.

### Consent for publication

It was clearly indicated on the information sheet provided to the study participants that data generated from the study will be anonymously communicated to the wider scientific community in the form of peer-reviewed scientific publication. Availability of data and materials: Data used to make the major conclusions of the study will be available together with this article and detailed data can be provided up on request. Competing interests: All authors declare that they don’t have competing interests. Funding: The study was supported by the Bill and Melinda Gates Foundation grant (INDIE OPP1173572) to FGT, TB and CD. The Armauer Hansen Research Institute (AHRI) has supported WC, TA, EH, and EG through its Core funding from Sida and Norad. PMI VectorLink Ethiopia Project for financial support of the adult collection. CLJ and TW were supported by Wellcome Trust /Royal Society fellowships awarded to TW (101285/Z/13/Z): http://www.wellcome.ac.uk; https://royalsociety.org

### Authors’ contributions

Conceived the study: HT, PM, SC, MM, MB, SI, CD, EG, TB, FGT; participated in guiding the field activities: PM, DD, SC, MM, MB, SI, JHK, AW, CD, EG, TB, FGT; collected the developmental stages (larvae and pupae): TA, EE; reared adult mosquitoes, collected blood samples, run feeding experiments, dissected mosquitoes: TA, EE, WC, SWB, DAM, EH, SKT, TT, AG, TT, TE, GY, SK, GS, SAS; conducted laboratory works: TA, HT, EE, LAM, WC, TW, SWB, KL, RH, CLJ, DAM, EH, SKT, TT, AG, TT, TE, FGT; analyzed data: TA, LAM, LMK, KS, TSC, SI, CD, EG, TB, FGT; drafted the manuscript: TA, HT, EE, LAM, TW, CLJ, CD, EG, TB, FGT; critically commented on the manuscript: JHK, AW, TSC, SI, CD, EG, TB, FGT. All authors read and approved the final manuscript.

## Acknowledgements

We would like to acknowledge the study participants for their willingness to donate blood samples and allow assessment of mosquito exposure. We thank the microscopists (Tewabech Lema and Tsehay Orlando) at Adama malaria clinic for their support. Appreciation also goes to the regional and district health officers for their collaboration and PMI VectorLink Ethiopia Project for training the field team. The drivers from AHRI played an important role on making the study successful. The findings and conclusions in this report are those of the author(s) and do not necessarily represent the official position of the Centers for Disease Control and Prevention.

