## Supplemental notes for "*Anopheles stephensi* as an emerging malaria vector in the Horn of Africa with high susceptibility to Ethiopian *Plasmodium vivax* and *Plasmodium falciparum* isolates"

**Aquatic habitat characterization**

A total of (n=85) potential larval habitats were surveyed in five days (May/June 2019) in Awash Sebat Kilo town using standardized check list/questionnaire (22 question sets) to capture important characteristics indicated in Table 1. Immatures were collected throughout the day. Altitude, latitude, longitude, sun light exposure, water turbidity, substrate type, presence of vegetation, predators and competitors were recorded for each site. The *Anopheles* larvae were separated from the culicine larvae and classified as early- ($1^{st}$, $2^{nd}$) or late-instar ($3^{rd}$ and $4^{th}$) stage and larval density was recorded by instars. The *Anopheles* larvae/pupae were transported to Adama malaria center with jars and transferred to larval tray for rearing to adult using the same but filtered water from the breeding site. **Table 1. Questionnaire for aquatic habitat characterization.**

| Number | Question | Response |
|---|---|---|
| 1 | Breeding site ID | |
| 2 | Survey Date | MM/DD/YY |
| 3 | Kebele | |
| 4 | Coordinates fo brreding habitat | |
| | Lat | |
| | Long | |
| | Altitude | |
| 5 | Where do the larval breeding habitat found? | 1 = Street |
| | | 2 = Public green area |
| | | 3 = Abandoned area (garbage area) |
| | | 4 = Public buildings (school, hospital) |
| | | 5 = Around household |
| | | 6 = Factory/brick production area |
| | | 7 = Other: (Specify)____________________ |
| 6 | Water body type / category | 1 = Permanent |
| | | 2 = Temporary |
| 7 | Water body type | 1 =Natural |
| | | 2= Manmade |
| 8 | Size of the breeding habitat | length*width*depth in meter |
| 9 | Shade status | 1 = Fully shaded |
| | | 2 = Partially shaded |
| 10 | Usage status of water container | 1 = in use |
| | | 2 = not in use |
| 11 | Purpose of water container | 1 = For household purpose |
| | | 2 = For Irrigation/gardening |
| | | 3 = For Animal drink |
| | | 4 = For constraction |
| | | 5 = Other (describe) ___________________ |
| 12 | What is the material that the contianer is made ? | 1 = Plastic |
| | | 2 = Metal |
| | | 3 = Cemented |
| | | 4 = Earth |
| | | 5 = other (describe) ___________________ |
| 13 | Water container type | 1 = Drum / Barrel |
| | | 2 = Steel Tank |
| | | 3 = Ceramic / cemented |
| | | 4 = Bucket |
| | | 5 = Tyer |
| | | 6 = Pots |
| | | 7 = Fiber jar |
| | | 8 = Other:(Specify) __________ |
| 14 | Is there predators | 1 = Yes |
| | | 2 = No |
| 15 | Inervention applied | 1 = Yes |
| | | 2 = No |
| 16 | Larvae present | 1 = Yes |
| | | 2 = No |
| 17 | If Q16 "Yes" measure the density | 1 = no larvae |
| | | 2 = < 10 |
| | | 3 = 10-50 |
| | | 4 = > 50 |
| 18 | Number of pupae in total dip | |
| 19 | Co-habitation by other spp or organism | 1 = Algae |
| | | 2 = Plants |
| | | 3 = Culex species |
| | | 4 = Aedes species |
| | | 5 = Decaying biological remnants |
| | | 6 = Other (specify) ____________________ |
| 20 | Description of larval habitat | 1 = clear |
| | | 2 = polluted |
| 21 | Water temperature of the breeding habitat | |
| 22 | Source of the water | Tap |
| | | Tap + rain |

Indicated in Table 2 is the outcome of the survey with each row indicating the aquatic habitats and the columns indicating the respective question numbers from the questionnaire. The responses are indicated with numbers in Table 2 referring to the specific code of the response in the alternatives given in the questionnaire (Table 1).

**Table 2. Data from the surveyed potential aquatic habitats**

| Question Study site | 1 | 2 | 3 | 4 | | | 5 | 6 | 7 | 8 | 9 | 10 | 11 | 12 | 13 | 14 | 15 | 16 | 17 | 18 | 19 | 20 | 21 | 22 |
|---|---|---|---|---|---|---|---|---|---|---|---|---|---|---|---|---|---|---|---|---|---|---|---|---|
| 1 | a001 | 31/05/2019 | 0 | 85908,1 | 400916,8 | 924 | 1 | 2 | 2 | 2*1,2*1,2 | 2 | 2 | NK | 2 | 2 | 2 | 2 | 1 | 2 | 0 | 4 | 2 | 26,5 | 1 |
| 2 | a002 | 31/05/2019 | 0 | 85912,5 | 400912,4 | 931 | 5 | 1 | 2 | 2*1*1 | 3 | 1 | 4 | 3 | 3 | 2 | 2 | 1 | 2 | 0 | 1 | 2 | 25 | 1 |
| 3 | a003 | 31/05/2019 | 0 | 85916,5 | 400912,2 | 928 | 5 | 1 | 2 | 2.5*2*2 | 3 | 1 | 4 | 3 | 3 | 2 | 2 | 1 | 2 | 0 | 0 | 1 | 25 | 1 |
| 4 | a004 | 31/05/2019 | 0 | 85917,1 | 400912,9 | 926 | 5 | 1 | 2 | 3*2*2 | 3 | 2 | NK | 3 | 3 | 2 | 2 | 2 | | 0 | 0 | 2 | 25,2 | 1 |
| 5 | a005 | 31/05/2019 | 0 | 85919,8 | 400931,6 | 925 | 5 | 1 | 2 | 2*1*1,5 | 3 | 1 | 4 | 3 | 3 | 2 | 2 | 1 | 2 | 0 | 1 | 1 | 24,5 | 1 |
| 6 | a006 | 31/05/2019 | 0 | 85925,6 | 400912,1 | 925 | 5 | 1 | 2 | 1.5*1*1 | 1 | 1 | 1 | 3 | 3 | 2 | 2 | 1 | 2 | 0 | 0 | 1 | 24,6 | 1 |
| 7 | a007 | 31/05/2019 | 0 | 85931,3 | 400920,9 | 926 | 5 | 2 | 2 | 2*1*1 | 2 | 1 | 4 | 2 | 2 | 2 | 2 | 1 | 2 | 0 | 0 | 1 | 27,5 | 1 |
| 8 | a008 | 31/05/2019 | 0 | 85932,2 | 400920,5 | 925 | 5 | 1 | 2 | 2*2*1 | 1 | 1 | 1 | 3 | 2 | 2 | 2 | 1 | 2 | 2 | 4 | 1 | 24,3 | 1 |
| 9 | a009 | 31/05/2019 | 0 | 85929,9 | 400919,3 | 923 | 5 | 1 | 2 | 2*1*1 | 2 | 1 | 1 | 3 | 3 | 2 | 2 | 1 | 2 | 0 | 0 | 1 | 24,5 | 1 |
| 10 | a010 | 31/05/2019 | 0 | 85928,7 | 400917,6 | 924 | 5 | 1 | 2 | 2.5*1.5*1.5 | 1 | 1 | 1 | 3 | 3 | 2 | 2 | 1 | 3 | 2 | 4 | 1 | 23 | 1 |
| 11 | a011 | 01/06/2019 | 0 | 85913,6 | 400915,4 | 921 | 5 | 1 | 2 | 2.5*1.5*1.5 | 3 | 1 | 4 | 3 | 3 | 2 | 2 | 1 | 4 | 0 | 0 | 1 | 26 | 1 |
| 12 | a012 | 01/06/2019 | 0 | 85918 | 400918,3 | 932 | 1 | 1 | 2 | 3*1.5*1.5 | 3 | 1 | 4 | 2 | 2 | 2 | 2 | 1 | 4 | 2 | 5 | 1 | 27,9 | 1 |
| 13 | a013 | 01/06/2019 | 0 | 85920,8 | 400917,1 | 927 | 5 | 1 | 2 | 2.5*1*1 | 1 | 1 | 4 | 3 | 3 | 2 | 2 | 1 | 2 | 13 | 1 | 1 | 25,5 | 1 |
| 14 | a014 | 01/06/2019 | 0 | 85924,7 | 400922,2 | 932 | 2 | 2 | 2 | 2*1*1 | 2 | 1 | 4 | 2 | 2 | 2 | 2 | 1 | 2 | 1 | 0 | 1 | 25,9 | 1 |
| 15 | a015 | 01/06/2019 | 0 | 85923,5 | 400922,7 | 929 | 5 | 2 | 2 | 1*1*1.5 | 2 | 1 | 1 | 2 | 2 | 2 | 2 | 1 | 2 | 0 | 4 | 1 | 26,2 | 1 |
| 16 | a016 | 01/06/2019 | 0 | 85924,7 | 400923,2 | 924 | 5 | 1 | 2 | 2.5*2.5*2.5 | 2 | 1 | 1 | 3 | 3 | 2 | 2 | 1 | 3 | 45 | 4 | 1 | 23,4 | 2 |
| 17 | a017 | 01/06/2019 | 0 | 85924,7 | 400923,7 | 931 | 2 | 2 | 2 | 1*1*1 | 2 | 1 | 1 | 2 | 2 | 2 | 2 | 1 | 3 | 5 | 11 | 1 | 25,5 | 2 |
| 18 | a018 | 01/06/2019 | 0 | 85924,5 | 400923,7 | 920 | 5 | 2 | 2 | 1*1*1 | 2 | 1 | 1 | 2 | 2 | 2 | 2 | 1 | 3 | 11 | 4 | 1 | 26,1 | 2 |
| 19 | a019 | 01/06/2019 | 0 | 85923,9 | 400923,5 | 927 | 5 | 2 | 2 | 0.9*0.9*1.2 | 1 | 1 | 1 | 1 | 7 | 2 | 2 | 2 | ND | 0 | 3 | 2 | 19,8 | 2 |
| 20 | a020 | 01/06/2019 | 0 | 85923,6 | 400924,8 | 924 | 5 | 2 | 2 | 0.9*0.9*1 | 1 | 1 | 1 | 1 | 7 | 2 | 2 | 1 | 2 | 1 | 9 | 1 | 21 | 1 |
| 21 | a021 | 01/06/2019 | 0 | 85924,6 | 400925 | 927 | 5 | 2 | 2 | 1*0.9*0.9 | 1 | 1 | 1 | 1 | 7 | 2 | 2 | 1 | 2 | 5 | 9 | 1 | 21 | 1 |
| 22 | a022 | 01/06/2019 | 0 | 85925,7 | 400925,7 | 925 | 6 | 1 | 2 | 2.5*2.5*2.5 | 2 | 2 | NK | 3 | 3 | 2 | 2 | 1 | 3 | 0 | 5 | 2 | 21,2 | 2 |
| 23 | a023 | 01/06/2019 | 0 | 85924,5 | 400929,3 | 924 | 5 | 1 | 2 | 4*3*1.5 | 3 | 1 | 1 | 3 | 2 | 2 | 2 | 1 | 2 | 0 | 8 | 2 | 23 | 1 |
| 24 | a024 | 01/06/2019 | 0 | 85857,9 | 400931,9 | 926 | 5 | 2 | 2 | 1.5*1.5*1 | 1 | 1 | 1 | 2 | 1 | 2 | 2 | 1 | 3 | 0 | 9 | 1 | 25 | 1 |
| 25 | a025 | 01/06/2019 | 0 | 85857,9 | 400932,1 | 930 | 5 | 1 | 2 | 2.5*1.5*1.5 | 1 | 2 | NK | 3 | 3 | 2 | 2 | 1 | 3 | 1 | 9 | 2 | 25,5 | 2 |
| 26 | a026 | 01/06/2019 | 0 | 85859,3 | 400932,1 | 925 | 6 | 1 | 2 | 3*2*1 | 1 | 1 | 4 | 3 | 3 | 2 | 2 | 1 | 3 | 0 | 0 | 1 | 24,6 | 1 |
| 27 | a027 | 01/06/2019 | 0 | 85859,3 | 400931,6 | 929 | 6 | 1 | 2 | 3*2*1.5 | 2 | 1 | 4 | 3 | 3 | 2 | 2 | 1 | 2 | 19 | 0 | 2 | 24 | 1 |
| 28 | a028 | 01/06/2019 | 0 | 85900,1 | 400931,7 | 930 | 5 | 1 | 2 | 2*2*1 | 3 | 2 | NK | 3 | 3 | 2 | 2 | 1 | 3 | 5 | 0 | 2 | 24,1 | 2 |
| 29 | a029 | 01/06/2019 | 0 | 85905,1 | 400932,5 | 925 | 6 | 1 | 2 | 4*4*1 | 3 | 1 | 4 | 3 | 3 | 2 | 2 | 1 | 3 | 0 | 1 | 1 | 23 | 1 |
| 30 | a030 | 01/06/2019 | 0 | 85904,8 | 400931,9 | 924 | 6 | 1 | 2 | 4*4*1 | 3 | 1 | 4 | 3 | 3 | 2 | 2 | 1 | 3 | 0 | 5 | 1 | 25 | 1 |
| 31 | a031 | 01/06/2019 | 0 | 85906 | 400932,3 | 923 | 6 | 1 | 2 | 8*4*2 | 3 | 1 | 4 | 3 | 3 | 2 | 2 | 1 | 3 | 3 | 5 | 1 | 25,5 | 1 |
| 32 | a032 | 01/06/2019 | 0 | 85908 | 400932,4 | 921 | 6 | 1 | 2 | 5*4*1 | 3 | 1 | 4 | 3 | 3 | 2 | 2 | 1 | 4 | 4 | 0 | 1 | 24 | 1 |
| 33 | a033 | 01/06/2019 | 0 | 85909,9 | 400932,6 | 921 | 6 | 1 | 2 | 4*3*1 | 3 | 1 | 4 | 3 | 3 | 2 | 2 | 1 | 4 | 13 | 8 | 2 | 25 | 1 |
| 34 | a034 | 01/06/2019 | 0 | 85916,9 | 400932,2 | 919 | 5 | 1 | 2 | 2.5*2*1 | 2 | 1 | 1 | 3 | 3 | 2 | 2 | 1 | 4 | 100 | 9 | 1 | 23 | 1 |
| 35 | a035 | 01/06/2019 | 0 | 85920,9 | 400934,5 | 917 | 5 | 2 | 2 | 1*1*1 | 1 | 1 | 1 | 1 | 7 | 2 | 2 | 1 | 2 | 0 | 0 | 1 | 23 | 1 |
| 36 | a036 | 01/06/2019 | 0 | 85924,7 | 400934,7 | 918 | 6 | 1 | 2 | 12*6*1*5 | 3 | 1 | 4 | 3 | 3 | 2 | 2 | 1 | 3 | 0 | 8 | 1 | 22 | 1 |
| 37 | a037 | 02/06/2019 | 0 | 85901,9 | 400933,4 | 936 | 6 | 1 | 2 | 3*2*1 | 1 | 2 | NK | 3 | 3 | 2 | 2 | 1 | 2 | 0 | 5 | 2 | 24 | 1 |
| 38 | a038 | 02/06/2019 | 0 | 85901,1 | 400934,1 | 934 | 5 | 2 | | 1*0.9*0.9 | 1 | 1 | 1 | 1 | 7 | 2 | 2 | 2 | 1 | 0 | 0 | 1 | 24 | 1 |
| 39 | a039 | 02/06/2019 | 0 | 85900,8 | 400934,1 | 932 | 5 | 2 | 2 | 1*0.9*0.9 | 1 | 1 | 1 | 2 | 1 | 2 | 2 | 2 | 1 | 0 | 9 | 1 | 23,4 | 2 |
| 40 | a040 | 02/06/2019 | 0 | 85901,3 | 400934,4 | 933 | 5 | 2 | 2 | 0.9*0.5*0.5 | 3 | 1 | 4 | 1 | 7 | 2 | 2 | 2 | 1 | 0 | 9 | 2 | 25 | 2 |
| 41 | a041 | 02/06/2019 | 0 | 85902,3 | 400935,1 | 935 | 5 | 1 | 2 | 3*1*1 | 2 | 1 | 1 | 2 | 2 | 2 | 2 | 1 | 2 | 2 | 0 | 1 | 25,5 | 1 |
| 42 | a042 | 02/06/2019 | 0 | 85903,7 | 400934,1 | 930 | 5 | 2 | 2 | 1*0.9*0.9 | 2 | 1 | 1 | 1 | 7 | 2 | 2 | 1 | 2 | 0 | 9 | 1 | 23 | 2 |
| 43 | a043 | 02/06/2019 | 0 | 85903,4 | 400934,5 | 934 | 5 | 2 | 2 | 0.4*0.4*0.4 | 1 | 1 | 1 | 2 | 1 | 2 | 2 | 1 | 2 | 0 | 4 | 1 | 25,4 | 1 |
| 44 | a044 | 02/06/2019 | 0 | 85903,7 | 400934,4 | 926 | 5 | 2 | 2 | 2*1*1 | 2 | 1 | 2 | 1 | 6 | 2 | 2 | 1 | 2 | 0 | 0 | 1 | 26,5 | 3 |
| 45 | a045 | 02/06/2019 | 0 | 85902,6 | 400934,6 | 932 | 7 | 2 | 2 | ND | 3 | 2 | NK | 1 | 5 | 2 | 2 | 1 | 2 | 0 | 4 | 1 | 25 | 2 |
| 46 | a046 | 02/06/2019 | 0 | 85902,6 | 400934,3 | 929 | 7 | 1 | 2 | 5*5*2 | 1 | 1 | 5 | 3 | 8 | 2 | 2 | 2 | 1 | 0 | 0 | 1 | 25 | 1 |
| 47 | a047 | 02/06/2019 | 0 | 85903,5 | 400933,7 | 935 | 5 | 2 | 2 | 1*0.9 | 1 | 1 | 1 | 1 | 7 | 2 | 2 | 2 | 1 | 0 | 3 | 1 | 25 | 3 |
| 48 | a048 | 02/06/2019 | 0 | 85903,7 | 400935,9 | 933 | 5 | 2 | 2 | 1*0.9 | 1 | 1 | 1 | 1 | 7 | 2 | 2 | 2 | 1 | 0 | 0 | 1 | 25 | 1 |
| 49 | a049 | 02/06/2019 | 0 | 85903,6 | 400938,4 | 933 | 5 | 2 | 2 | ND | 3 | 2 | NK | 1 | 5 | 2 | 2 | 2 | 1 | 0 | 0 | 2 | 26,5 | 2 |
| 50 | a050 | 02/06/2019 | 0 | 85905,4 | 400940,7 | 927 | 6 | 2 | 2 | ND | 3 | 2 | NK | 1 | 5 | 2 | 2 | 1 | 2 | 1 | 9 | 2 | 25,5 | 2 |
| 51 | a051 | 02/06/2019 | 0 | 85906,8 | 400940,9 | 926 | 5 | 1 | 2 | 2*2*1 | 3 | 2 | NK | 3 | 3 | 2 | 2 | 1 | 2 | 0 | 0 | 2 | 25 | 1 |
| 52 | a052 | 02/06/2019 | 0 | 85908,5 | 400941,4 | 920 | 5 | 2 | 2 | ND | 3 | 2 | NK | 1 | 5 | 2 | 2 | 2 | 1 | 0 | 0 | 2 | 25 | 2 |
| 53 | a053 | 02/06/2019 | 0 | 85908,8 | 400943,7 | 925 | 1 | 2 | 2 | ND | 3 | 2 | NK | 1 | 5 | 2 | 2 | 2 | 1 | 0 | 0 | 2 | 25 | 2 |
| 54 | a054 | 02/06/2019 | 0 | 85911,4 | 400943,7 | 925 | 5 | 2 | 2 | ND | 3 | 1 | 2 | 1 | 6 | 2 | 2 | 2 | 1 | 0 | 0 | 2 | 26 | 2 |
| 55 | a055 | 02/06/2019 | 0 | 85911 | 400944,2 | 926 | 5 | 2 | 2 | 1*0.9 | 3 | 1 | 1 | 1 | 7 | 2 | 2 | 2 | 1 | 0 | 4 | 1 | 25 | 1 |
| 56 | a056 | 02/06/2019 | 0 | 85913,2 | 400945,7 | 928 | 7 | 2 | 2 | 1*0.9 | 3 | 1 | 4 | 2 | 1 | 2 | 2 | 1 | 2 | 0 | 0 | 1 | 24 | 1 |
| 57 | a057 | 02/06/2019 | 0 | 85907,6 | 400939,9 | 931 | 5 | 2 | 2 | 1*0.9 | 1 | 1 | 1 | 1 | 7 | 2 | 2 | 2 | 1 | 0 | 0 | 1 | 25 | 1 |
| 58 | a058 | 02/06/2019 | 0 | 85908,3 | 400938,8 | 919 | 5 | 2 | 2 | 1*0.9 | 3 | 1 | 1 | 1 | 7 | 2 | 2 | 2 | 1 | 0 | 0 | 1 | 24,5 | 1 |
| 59 | a059 | 02/06/2019 | 0 | 85910,7 | 400939,1 | 922 | 5 | 2 | 2 | 0.9*0.5 | 1 | 1 | 1 | 2 | 1 | 2 | 2 | 1 | 2 | 0 | 9 | 1 | 25 | 1 |
| 60 | a060 | 02/06/2019 | 0 | 85910,7 | 400939,1 | 927 | 5 | 2 | 2 | 1*0.5 | 2 | 1 | 1 | 1 | 7 | 2 | 2 | 2 | 1 | 0 | 4 | 1 | 24 | 2 |
| 61 | a061 | 08/06/2019 | 2 | 90024,2 | 401041,7 | 901 | 1 | 1 | 2 | 4*4*1 | 3 | 2 | NK | 3 | 3 | 2 | 2 | 1 | 4 | 7 | 6 | 1 | 23,3 | 1 |
| 62 | a062 | 08/06/2019 | 2 | 90023 | 401041,4 | 899 | 6 | 1 | 2 | 1*0.5*0.5 | 3 | 1 | 4 | 3 | 3 | 2 | 2 | 1 | 3 | 22 | 6 | 1 | 25 | 1 |
| 63 | a063 | 08/06/2019 | 2 | 90024,1 | 401042,4 | 898 | 6 | 1 | 2 | 4*4*2 | 3 | 1 | 4 | 3 | 3 | 2 | 2 | 1 | 4 | 20 | 1 | 2 | 28,5 | 1 |
| 64 | a064 | 08/06/2019 | 2 | 90023,7 | 401044,5 | 897 | 6 | 1 | 2 | 6*2*2 | 3 | 1 | 4 | 3 | 3 | 2 | 2 | 1 | 3 | 0 | 1 | 1 | 28 | 1 |
| 65 | a065 | 08/06/2019 | 2 | 90028,6 | 401042,5 | 901 | 6 | 2 | 2 | 3*3*1 | 2 | 1 | 4 | 2 | 2 | 2 | 2 | 1 | 2 | 0 | 0 | 1 | 27 | 1 |
| 66 | a066 | 08/06/2019 | 2 | 90023,1 | 401038,5 | 902 | 6 | 1 | 2 | 4*3*2 | 3 | 1 | 4 | 3 | 3 | 2 | 2 | 1 | 3 | 0 | 0 | 1 | 26,5 | 1 |
| 67 | a067 | 08/06/2019 | 2 | 90019,1 | 401035,4 | 900 | 6 | 1 | 2 | 4*3*1 | 3 | 1 | 4 | 3 | 3 | 2 | 2 | 1 | 4 | 13 | 0 | 1 | 27 | 1 |
| 68 | a068 | 08/06/2019 | 2 | 9007,4 | 401026,1 | 909 | 5 | 2 | 2 | ND | 3 | 1 | 1 | 1 | 5 | 2 | 2 | 1 | 4 | 15 | 7 | 2 | 28 | 1 |
| 69 | a069 | 08/06/2019 | 1 | 85853 | 400940,5 | 935 | 6 | 1 | 2 | 3*1*2 | 3 | 1 | 4 | 3 | 3 | 2 | 2 | 1 | 2 | 0 | 0 | 1 | 27 | 1 |
| 70 | a070 | 08/06/2019 | 1 | 85856,8 | 400936,5 | 926 | 6 | 1 | 2 | 2*2*1 | 3 | 1 | 4 | 3 | 3 | 2 | 2 | 2 | 1 | 0 | 1 | 2 | 26 | 1 |
| 71 | a071 | 08/06/2019 | 1 | 85857,5 | 400936 | 930 | 6 | 1 | 2 | 2*2*1 | 3 | 1 | 4 | 3 | 3 | 2 | 2 | 2 | 1 | 0 | 1 | NK | 25 | 2 |
| 72 | a072 | 08/06/2019 | 1 | 85900,4 | 400939,3 | 928 | 6 | 1 | 2 | 4*4*2 | 3 | 1 | 4 | 3 | 3 | 2 | 2 | 1 | 2 | 0 | 1 | 2 | 26 | 1 |
| 73 | a073 | 08/06/2019 | 1 | 85900,3 | 400939 | 930 | 6 | 1 | 2 | 3*1*1 | 2 | 1 | 5 | 2 | 2 | 2 | 2 | 1 | 3 | 0 | 9 | 1 | 26 | 1 |
| 74 | a074 | 08/06/2019 | 1 | 85901,1 | 400940 | 928 | 6 | 1 | 2 | 6*3*2 | 1 | 1 | 4 | 3 | 3 | 2 | 2 | 1 | 2 | 0 | 0 | 1 | 25 | 1 |
| 75 | a075 | 08/06/2019 | 1 | 85901,1 | 400940,8 | 923 | 6 | 1 | 2 | 4*4*2 | 2 | 1 | 4 | 3 | 3 | 2 | 2 | 1 | 2 | 0 | 8 | 2 | 27 | 1 |
| 76 | a076 | 08/06/2019 | 1 | 85901,6 | 400941,4 | 926 | 5 | 2 | 2 | ND | 3 | 2 | NK | 1 | 5 | 2 | 2 | 1 | 2 | 0 | 4 | 2 | 23 | 2 |
| 77 | a077 | 08/06/2019 | 1 | 85902,1 | 400941,2 | 923 | 5 | 2 | 2 | 2*2*1 | 2 | 1 | 1 | 2 | 2 | 2 | 2 | 1 | 2 | 0 | 0 | 1 | 24,5 | 1 |
| 78 | a078 | 08/06/2019 | 1 | 85904 | 400942 | 925 | 6 | 1 | 2 | 4*4*2 | 3 | 1 | 4 | 3 | 3 | 2 | 2 | 2 | 1 | 0 | 0 | 1 | 27 | 1 |
| 79 | a082 | 09/06/2019 | 1 | 85904,8 | 400942,9 | 924 | 5 | 1 | 2 | 1*1*1.8 | 2 | 1 | 1 | 3 | 3 | 2 | 2 | 1 | 2 | 0 | 9 | 2 | 25,5 | 1 |
| 80 | a083 | 09/06/2019 | 1 | 85904,9 | 400942,7 | 925 | 5 | 2 | 2 | 1*1*1.5 | 2 | 1 | 1 | 1 | 7 | 2 | 2 | 1 | 3 | 0 | 0 | 1 | 26 | 1 |
| 81 | a084 | 09/06/2019 | 1 | 85905,4 | 400942,4 | 928 | 1 | 1 | 2 | 1*1*1.5 | 1 | 1 | 1 | 3 | 3 | 2 | 2 | 2 | 1 | 0 | 4 | 1 | 25 | 1 |
| 82 | a085 | 09/06/2019 | 1 | 85907,7 | 400945 | 929 | 5 | 2 | 2 | 1*0.5*0.5 | 2 | 1 | 2 | 2 | 2 | 2 | 2 | 1 | 3 | 0 | 9 | 1 | 27 | 1 |
| 83 | a086 | 09/06/2019 | 1 | 85910,6 | 400946,1 | 930 | 6 | 1 | 2 | 4*4*1 | 3 | 1 | 4 | 3 | 3 | 2 | 2 | 1 | 2 | 0 | 1 | 2 | 26,5 | 1 |
| 84 | a087 | 09/06/2019 | 1 | 85910 | 400946,4 | 928 | 6 | 1 | 2 | 4*4*1 | 2 | 1 | 4 | 3 | 3 | 2 | 2 | 2 | 1 | 0 | 1 | 2 | 26 | 1 |
| 85 | a088 | 09/06/2019 | 1 | 85910,7 | 400946,8 | 926 | 6 | 1 | 2 | 5*4*2 | 2 | 1 | 4 | 3 | 3 | 2 | 2 | 1 | 3 | 0 | 10 | 2 | 25 | 1 |

*ND, not done; NK, not known; values in the third column of question 4 refer to altitude measured above sea level; values in question 8 refer to the dimension of the aquatic habitat in meters*

Number of larvae/pupae collected in each of the 10 dips (indicated in columns) is shown for each surveyed habitat (indicated in rows) in Table 3.

**Table 3. Larval density/dip/aquatic habitat from Awash Sebat kilo for feeding assay**

| code | dip1 | dip2 | dip3 | dip4 | dip5 | dip6 | dip7 | dip8 | dip9 | dip10 | diptotal |
|---|---|---|---|---|---|---|---|---|---|---|---|
| a001 | 3 | 3 | 2 | 2 | 2 | 3 | 1 | 1 | 2 | 3 | 22 |
| a002 | 8 | 2 | 6 | 9 | 4 | 12 | 7 | 14 | 23 | 5 | 90 |
| a003 | 0 | 0 | 0 | 1 | 4 | 2 | 3 | 2 | 16 | 3 | 31 |
| a005 | 1 | 7 | 1 | 0 | 0 | 1 | 0 | 8 | 5 | 7 | 30 |
| a006 | 6 | 3 | 4 | 10 | 2 | 1 | 7 | 7 | 0 | 11 | 51 |
| a007 | 2 | 4 | 2 | 0 | 6 | 3 | 3 | 8 | 8 | 11 | 47 |
| a008 | 0 | 1 | 0 | 1 | 2 | 0 | 1 | 2 | 3 | 1 | 11 |
| a009 | 3 | 0 | 5 | 0 | 0 | 0 | 1 | 1 | 0 | 0 | 10 |
| a010 | 9 | 10 | 21 | 3 | 6 | 7 | 7 | 4 | 19 | 35 | 121 |
| a011 | 50 | 70 | 20 | 35 | 45 | 25 | 65 | 50 | 40 | 30 | 430 |
| a012 | 70 | 25 | 80 | 50 | 130 | 80 | 120 | 90 | 60 | 80 | 785 |
| a013 | 16 | 8 | 6 | 12 | 4 | 5 | 9 | 11 | 3 | 5 | 79 |
| a014 | 1 | 5 | 0 | 1 | 0 | 0 | 2 | 1 | 0 | 3 | 13 |
| a015 | 1 | 0 | 1 | 0 | 0 | 2 | 1 | 0 | 1 | 1 | 7 |
| a016 | 30 | 25 | 45 | 20 | 50 | 25 | 10 | 20 | 30 | 40 | 295 |
| a017 | 14 | 8 | 6 | 4 | 12 | 5 | 15 | 12 | 6 | 18 | 100 |
| a018 | 30 | 25 | 15 | 20 | 30 | 20 | 23 | 14 | 32 | 16 | 225 |
| a020 | 1 | 0 | 0 | 5 | 10 | 6 | 9 | 4 | 3 | 2 | 40 |
| a021 | 4 | 0 | 2 | 3 | 0 | 4 | 1 | 0 | 1 | 1 | 16 |
| a022 | 15 | 20 | 10 | 12 | 25 | 20 | 35 | 70 | 65 | 40 | 312 |
| a023 | 3 | 4 | 2 | 5 | 6 | 1 | 2 | 4 | 1 | 7 | 35 |
| a024 | 35 | 20 | 15 | 30 | 30 | 35 | 25 | 20 | 30 | 40 | 280 |
| a025 | 21 | 26 | 75 | 17 | 24 | 16 | 11 | 32 | 40 | 14 | 276 |
| a026 | 4 | 15 | 6 | 4 | 8 | 6 | 11 | 17 | 4 | 7 | 82 |
| a027 | 11 | 15 | 0 | 2 | 5 | 4 | 12 | 7 | 3 | 9 | 68 |
| a028 | 35 | 45 | 40 | 55 | 30 | 30 | 35 | 45 | 50 | 40 | 405 |
| a029 | 10 | 15 | 20 | 25 | 35 | 17 | 12 | 25 | 20 | 15 | 194 |
| a030 | 22 | 18 | 15 | 26 | 20 | 18 | 24 | 10 | 25 | 17 | 195 |
| a031 | 6 | 17 | 10 | 12 | 14 | 8 | 16 | 25 | 34 | 18 | 160 |
| a032 | 50 | 55 | 45 | 40 | 60 | 35 | 70 | 50 | 30 | 40 | 475 |
| a033 | 75 | 70 | 80 | 100 | 90 | 70 | 75 | 80 | 50 | 80 | 770 |
| a034 | 42 | 68 | 36 | 25 | 30 | 66 | 78 | 35 | 45 | 40 | 465 |
| a035 | 5 | 6 | 4 | 10 | 5 | 3 | 11 | 8 | 7 | 0 | 59 |
| a036 | 40 | 35 | 40 | 30 | 40 | 55 | 25 | 30 | 25 | 35 | 355 |
| a037 | 3 | 10 | 15 | 9 | 7 | 4 | 6 | 4 | 12 | 15 | 85 |
| a041 | 1 | 0 | 3 | 5 | 2 | 0 | 6 | 7 | 11 | 1 | 36 |
| a042 | 1 | 4 | 6 | 3 | 5 | 3 | 1 | 0 | 6 | 4 | 33 |
| a043 | 3 | 4 | 1 | 0 | 4 | 3 | 6 | 7 | 1 | 2 | 31 |
| a044 | 1 | 5 | 1 | 0 | 3 | 4 | 3 | 2 | 1 | 0 | 20 |
| a045 | 1 | 0 | 2 | 1 | 0 | 0 | 1 | 0 | 1 | 0 | 6 |
| a050 | 1 | 0 | 1 | 0 | 1 | 0 | 0 | 2 | 0 | 1 | 6 |
| a051 | 5 | 4 | 5 | 3 | 9 | 4 | 1 | 0 | 1 | 2 | 34 |
| a056 | 5 | 6 | 1 | 5 | 2 | 9 | 7 | 1 | 1 | 0 | 37 |
| a059 | 3 | 2 | 3 | 1 | 2 | 4 | 1 | 0 | 6 | 2 | 24 |
| a061 | 50 | 60 | 55 | 75 | 25 | 35 | 60 | 80 | 30 | 50 | 520 |
| a062 | 19 | 25 | 30 | 14 | 9 | 20 | 30 | 15 | 45 | 55 | 262 |
| a063 | 75 | 30 | 45 | 40 | 25 | 20 | 40 | 70 | 19 | 60 | 424 |
| a064 | 25 | 15 | 30 | 10 | 35 | 30 | 20 | 70 | 15 | 40 | 290 |
| a065 | 14 | 20 | 2 | 4 | 2 | 12 | 7 | 1 | 4 | 3 | 69 |
| a066 | 4 | 12 | 6 | 4 | 2 | 3 | 17 | 11 | 21 | 2 | 82 |
| a067 | 45 | 70 | 40 | 50 | 60 | 35 | 25 | 20 | 50 | 70 | 465 |
| a068 | 40 | 30 | 50 | 65 | 50 | 75 | 70 | 60 | 40 | 65 | 545 |
| a069 | 7 | 5 | 4 | 6 | 4 | 3 | 0 | 1 | 11 | 5 | 46 |
| a072 | 3 | 4 | 2 | 1 | 6 | 9 | 2 | 4 | 1 | 7 | 39 |
| a073 | 25 | 10 | 30 | 45 | 15 | 20 | 10 | 10 | 15 | 24 | 204 |
| a074 | 2 | 6 | 1 | 4 | 3 | 1 | 0 | 5 | 3 | 9 | 34 |
| a075 | 5 | 1 | 2 | 4 | 1 | 3 | 0 | 9 | 6 | 8 | 39 |
| a076 | 4 | 3 | 2 | 7 | 10 | 5 | 11 | 8 | 13 | 4 | 67 |
| a077 | 0 | 0 | 1 | 2 | 0 | 1 | 3 | 0 | 4 | 2 | 13 |
| a082 | 4 | 5 | 3 | 2 | 1 | 4 | 3 | 5 | 6 | 7 | 40 |
| a083 | 25 | 20 | 30 | 15 | 9 | 30 | 40 | 10 | 19 | 15 | 213 |
| a085 | 35 | 40 | 25 | 40 | 50 | 30 | 45 | 40 | 25 | 20 | 350 |
| a086 | 2 | 1 | 6 | 5 | 4 | 2 | 1 | 0 | 5 | 2 | 28 |
| a088 | 12 | 17 | 19 | 9 | 6 | 11 | 16 | 22 | 14 | 28 | 154 |

**Adult mosquitoes resting, biting and host preference behavior and egg-ridge study**

Resting, feeding and host preference behavior of *An. stephensi* was assessed using five entomological sampling techniques: i) Centers for Disease Control (CDC) light trap, ii) human landing catches (HLC), iii) pyrethrum spray sheet collection (PSC), iv) aspiration from animal shelters, and v) cattle baited traps. The CDC light traps (Model 512; John W. Hock Company, Gainesville, FL, USA) were set 1 meter above the ground on a wall or roof, both indoors and outdoors on 15 randomly selected households for two nights that makes a total of 60 traps in 30 nights. Indoor traps were hanged at the foot edge of the person who slept under untreated bed net (Lines, Curtis et al. 1991). Other occupants in the houses were left to use LLINs provided by the control program as part of the routine malaria control. The traps were switched on at 6:00PM and off at 6:00AM the next morning in each sampling night. PSC were conducted from 6:00AM to 2:00AM on five randomly selected households' per-day and 20 households were included in each round of sampling, thus a total of 60 households were sampled in three rounds. The HLC were conducted in nine selected households both in and outdoors that was repeated the next day. Locally trained entomology technicians were employed to collect female *Anopheles* by standard mouth aspirator from 6:00PM to 6:00AM from both indoors and outdoors. Two collectors were assigned at a time for each house (one outdoors and one indoors) in shifts of 6 hours (the first shift being 6:00PM – 12:00PM and the second from 12:00PM to 6:00AM). Collectors in the same shift swop each other between outdoors and indoors every hour after recording their findings on the checklist to avoid bias due to individual variation in attraction and competence. In addition to this, animal sheds were inspected using HC and cattle bait trap were conducted for collecting mosquitos biting and resting in animal shelters. Mosquitoes resting in animal shelters and cattle bait traps were collected using standard mouth aspirator for 30 minutes in each, from 5:30AM to 6:00AM. All collected *Anopheles* mosquitoes were counted and sorted out morphologically to species level (Gillies and Coetee 1987, Das, Rajagopal et al. 1990) and by their abdominal stage into unfed, freshly fed, half-gravid or gravid (Mobilization, Team et al. 2003), except those collected by HLC. In animal shelter with high number of mosquito collection was repeated the next morning.

Survey results for different adult catch methods is indicated in Table 4 for each of the three localities (Kebele) in Awash Sebat Kilo town with the following representations in column 'village': 1 = Alalamo; 2=Lemlefan; 3=Kedaba. The methodology refers to 1=CDC light trap; 2=Hand Collection; 3=Human Landing Catch. The column 'inout' refers to if the mosquito was caught indoor (0), outdoors (1) or tent (2). Species refers to *An. gambiae* (0), *An. stephensi* (1), or *An. pharoensis* (2). The feeding status (abdominal stage) of each of the mosquito is indicated in the last column: unfed (0), fed (1), gravid (2), or half gravid (3).

### Table 4. Adult *An. stephensi* surveillance data from Awash Sebat Kilo town

| sn | housen | idsample | date | village | method | inout | species | abdomenstage |
|---|---|---|---|---|---|---|---|---|
| 1 | HH01 | G01 | 28/08/2019 | 1 | 1 | 0 | 0 | 1 |
| 2 | HH01 | G02 | 28/08/2020 | 1 | 1 | 0 | 0 | 1 |
| 3 | HH01 | G03 | 28/08/2021 | 1 | 1 | 0 | 0 | 1 |
| 4 | HH01 | S01 | 28/08/2022 | 1 | 1 | 0 | 1 | 0 |
| 5 | HH01 | S02 | 28/08/2023 | 1 | 1 | 0 | 1 | 1 |
| 6 | HH01 | S03 | 28/08/2024 | 1 | 1 | 0 | 1 | 1 |
| 7 | HH01 | S04 | 28/08/2025 | 1 | 1 | 0 | 1 | 1 |
| 8 | HH01 | S05 | 28/08/2026 | 1 | 1 | 0 | 1 | 1 |
| 9 | HH01 | S06 | 28/08/2027 | 1 | 1 | 0 | 1 | 2 |
| 10 | HH01 | S07 | 28/08/2028 | 1 | 1 | 0 | 1 | 0 |
| 11 | HH01 | S08 | 28/08/2029 | 1 | 1 | 0 | 1 | 1 |
| 12 | HH02 | S09 | 29/08/2019 | 1 | 1 | 0 | 1 | 2 |
| 13 | HH02 | S10 | 29/08/2019 | 1 | 1 | 0 | 1 | 0 |
| 14 | HH02 | S11 | 29/08/2019 | 1 | 1 | 0 | 1 | 1 |
| 15 | HH02 | S12 | 29/08/2019 | 1 | 1 | 0 | 1 | 1 |
| 16 | HH02 | S13 | 29/08/2019 | 1 | 1 | 0 | 1 | 0 |
| 17 | HH02 | S14 | 29/08/2019 | 1 | 1 | 0 | 1 | 1 |
| 18 | HH02 | S15 | 29/08/2019 | 1 | 1 | 0 | 1 | 1 |
| 19 | HH02 | S16 | 29/08/2019 | 1 | 1 | 0 | 1 | 3 |
| 20 | HH02 | S17 | 29/08/2019 | 1 | 1 | 0 | 1 | 1 |
| 21 | HH03 | S18 | 30/08/2019 | 1 | 1 | 0 | 1 | 1 |
| 22 | HH03 | S19 | 30/08/2019 | 1 | 1 | 0 | 1 | |
| 23 | HH03 | S20 | 30/08/2019 | 1 | 1 | 0 | 1 | 1 |
| 24 | HH03 | G04 | 30/08/2019 | 1 | 1 | 0 | 0 | 1 |
| 25 | HH03 | G05 | 30/08/2019 | 1 | 1 | 0 | 0 | 1 |
| 26 | HH03 | G06 | 30/08/2019 | 1 | 1 | 0 | 0 | 1 |
| 27 | HH04 | G07 | 28/08/2019 | 1 | 2 | 1 | 0 | 1 |
| 28 | HH04 | S21 | 28/08/2019 | 1 | 2 | 1 | 1 | 1 |
| 29 | HH04 | S22 | 28/08/2019 | 1 | 2 | 1 | 1 | 1 |
| 30 | HH04 | S23 | 28/08/2019 | 1 | 2 | 1 | 1 | 1 |
| 31 | HH04 | S24 | 28/08/2019 | 1 | 2 | 1 | 1 | 1 |
| 32 | an1 | S25 | 22/09/2019 | 2 | 1 | 0 | 1 | 1 |
| 33 | tenet-d1 | S26 | 22/09/2019 | 3 | 1 | 0 | 1 | 1 |
| 34 | tenet-d1 | S27 | 22/09/2019 | 3 | 1 | 0 | 1 | 1 |
| 35 | hh1-d1 | G08 | 22/09/2019 | 2 | 0 | 0 | 0 | 1 |
| 36 | hh2-d1 | G09 | 22/09/2019 | 2 | 0 | 0 | 0 | 0 |
| 37 | hh2-d1 | G10 | 22/09/2019 | 2 | 0 | 0 | 0 | 1 |
| 38 | hh2-d1 | G11 | 22/09/2019 | 2 | 0 | 0 | 0 | 1 |
| 39 | hh2-d1 | G12 | 22/09/2019 | 2 | 0 | 0 | 0 | 1 |
| 40 | hh2-d1 | G13 | 22/09/2019 | 2 | 0 | 0 | 0 | 2 |
| 41 | tenet-d2 | S28 | 23/09/2019 | 3 | 1 | 2 | 1 | 2 |
| 42 | tenet-d2 | S29 | 23/09/2019 | 3 | 1 | 2 | 1 | 1 |
| 43 | an2 | S30 | 23/09/2019 | 2 | 1 | 0 | 1 | 1 |
| 44 | hh2-d2 | G14 | 23/09/2019 | 2 | 0 | 0 | 0 | 0 |
| 45 | hh1-d2 | S31 | 23/09/2019 | 2 | 2 | 1 | 1 | 0 |
| 46 | hh1-d2 | G15 | 23/09/2019 | 2 | 2 | 0 | 0 | 1 |
| 47 | an-3 | S32 | 23/09/2019 | 1 | 1 | 0 | 1 | 1 |
| 48 | an-3 | S33 | 23/09/2019 | 1 | 1 | 0 | 1 | 1 |
| 49 | an-3 | S34 | 23/09/2019 | 1 | 1 | 0 | 1 | 1 |
| 50 | an-3 | S35 | 23/09/2019 | 1 | 1 | 0 | 1 | 1 |
| 51 | an-3 | S36 | 23/09/2019 | 1 | 1 | 0 | 1 | 1 |
| 52 | an-3 | S37 | 23/09/2019 | 1 | 1 | 0 | 1 | 0 |
| 53 | an-3 | S38 | 23/09/2019 | 1 | 1 | 0 | 1 | 2 |
| 54 | an-3 | S39 | 23/09/2019 | 1 | 1 | 0 | 1 | 0 |
| 55 | an-3 | S40 | 23/09/2019 | 1 | 1 | 0 | 1 | 1 |
| 56 | an-3 | S41 | 23/09/2019 | 1 | 1 | 0 | 1 | 1 |
| 57 | an-3 | S42 | 23/09/2019 | 1 | 1 | 0 | 1 | 1 |
| 58 | an-3 | S43 | 23/09/2019 | 1 | 1 | 0 | 1 | 1 |
| 59 | an-3 | S44 | 23/09/2019 | 1 | 1 | 0 | 1 | 1 |
| 60 | an-3 | S45 | 23/09/2019 | 1 | 1 | 0 | 1 | 1 |
| 61 | an-3 | S46 | 23/09/2019 | 1 | 1 | 0 | 1 | 1 |
| 62 | an-3 | S47 | 23/09/2019 | 1 | 1 | 0 | 1 | 1 |
| 63 | an-3 | S48 | 23/09/2019 | 1 | 1 | 0 | 1 | 1 |
| 64 | an-3 | G16 | 23/09/2019 | 1 | 1 | 0 | 0 | 1 |
| 65 | an-4 | S49 | 24/09/2019 | 1 | 1 | 0 | 1 | 1 |
| 66 | an-4 | S50 | 24/09/2019 | 1 | 1 | 0 | 1 | 1 |
| 67 | an-4 | S51 | 24/09/2019 | 1 | 1 | 0 | 1 | 1 |
| 68 | an-4 | S52 | 24/09/2019 | 1 | 1 | 0 | 1 | 1 |
| 69 | an-4 | S53 | 24/09/2019 | 1 | 1 | 0 | 1 | 1 |
| 70 | an-4 | S54 | 24/09/2019 | 1 | 1 | 0 | 1 | 1 |
| 71 | an-5 | S55 | 24/09/2019 | 1 | 1 | 0 | 1 | 1 |
| 72 | an-5 | S56 | 24/09/2019 | 1 | 1 | 0 | 1 | 1 |
| 73 | an-5 | S57 | 24/09/2019 | 1 | 1 | 0 | 1 | 1 |
| 74 | an-5 | S58 | 24/09/2019 | 1 | 1 | 0 | 1 | 1 |
| 75 | an-5 | S59 | 24/09/2019 | 1 | 1 | 0 | 1 | 1 |
| 76 | an-5 | S60 | 24/09/2019 | 1 | 1 | 0 | 1 | 1 |
| 77 | tenet-3 | S61 | 24/09/2019 | 3 | 1 | 2 | 1 | 1 |
| 78 | hh3 | S62 | 25/09/2019 | 2 | 2 | 1 | 1 | 1 |
| 79 | tenet-4 | S63 | 26/09/2019 | 3 | 1 | 2 | 1 | 1 |
| 80 | tenet-4 | S64 | 26/09/2019 | 3 | 1 | 2 | 1 | 1 |
| 81 | tenet-4 | S65 | 26/09/2019 | 3 | 1 | 2 | 1 | 1 |
| 82 | tenet-4 | S66 | 26/09/2019 | 3 | 1 | 2 | 1 | 1 |
| 83 | tenet-4 | S67 | 26/09/2019 | 3 | 1 | 2 | 1 | 1 |
| 84 | an-6 | S68 | 26/09/2019 | 2 | 1 | 0 | 1 | 1 |
| 85 | an-6 | S69 | 26/09/2019 | 2 | 1 | 0 | 1 | 1 |
| 86 | an-6 | S70 | 26/09/2019 | 2 | 1 | 0 | 1 | 1 |
| 87 | an-6 | S71 | 26/09/2019 | 2 | 1 | 0 | 1 | 1 |
| 88 | an-6 | S72 | 26/09/2019 | 2 | 1 | 0 | 1 | 1 |
| 89 | an-6 | p01 | 26/09/2019 | 2 | 1 | 0 | 2 | 1 |

Optimum starvation time for the *An. stephensi* raised from wild collected larvae and pupae was assessed in three different experiments with 2, 3, 4 or 5 hours before feeding evaluated in each experiment (Table 5).

**Table 5. Feeding optimization for *An. stephensi* raised from wild collected larvae/pupae**

| Code of Experiment | Starvation hour prior feeding | Number of fed mosquitoes | Feeding efficiency, % of blood fed |
|---|---|---|---|
| E01-01 | 5 | 58 | 39.2 |
| E01-02 | 4 | 68 | 45.9 |
| E01-03 | 3 | 85 | 53.8 |
| E01-04 | 2 | 86 | 50.3 |
| E02-01 | 5 | 24 | 36.4 |
| E02-02 | 4 | 31 | 42.5 |
| E02-03 | 3 | 29 | 44.6 |
| E02-04 | 2 | 17 | 28.3 |
| E03-01 | 5 | 33 | 36.2 |
| E03-02 | 4 | 46 | 48.9 |
| E03-03 | 3 | 44 | 51.6 |
| E03-04 | 2 | 35 | 35.0 |

### Egg morphology identification

Fully fed mosquitoes were identified morphologically and *An. stephensi* were kept in mini paper cups at the Adama field laboratory at ambient conditions and were allowed to lay eggs on filter papers soaked in water on a cotton roll. A total of 10 eggs per female were collected and the egg-float ridge numbers were counted using Stereo-Microscope at 10X magnification and categorized based on criteria for biological forms: type, intermediate, and mysorensis (Subbarao, Vasantha et al. 1987). All three forms of *An. stephensi*, i.e. type-form, intermediate and mysorensis were observed in urban areas (Subbarao, Vasantha et al. 1987) with the 'type' occurring only in urban areas whereas only the mysorensis and intermediate forms are seen in rural areas (Oshaghi, Yaaghoobi et al. 2006, Mehravaran, Vatandoost et al. 2012, Nagpal, Srivastava et al. 2012). All *Anopheles* were preserved individually in tube containing self-indicating silica gel desiccant beads (Geejay Chemicals Ltd) covered with cotton pads and kept at -20$^{o}$C until further use. As indicated in Table 6, the median number of ridges among the eggs laid by individual females was 15 (interquartile range, 14-16; range 13-21).

**Table 6. Egg morphology of *An. stephensi***

| code | morphological ID | number of eggs laid | egg1 | egg2 | egg3 | egg4 | egg5 | egg6 | egg7 | egg8 | egg9 | egg10 |
|---|---|---|---|---|---|---|---|---|---|---|---|---|
| m1 | *An. stephensi* | 10 | 13 | 13 | 13 | 12 | 13 | 13 | 13 | 14 | 14 | 13 |
| m2 | *An. stephensi* | 65 | 17 | 19 | 18 | 19 | 16 | 15 | 18 | 14 | 16 | 13 |
| m3 | *An. stephensi* | 74 | 16 | 15 | 14 | 17 | 17 | 18 | 19 | 16 | 17 | 14 |
| m4 | *An. stephensi* | 55 | 14 | 17 | 14 | 16 | 16 | 19 | 18 | 17 | 16 | 16 |
| m5 | *An. stephensi* | 100 | 13 | 14 | 14 | 20 | 21 | 16 | 18 | 17 | 16 | 16 |
| m6 | *An. stephensi* | 84 | 14 | 16 | 14 | 22 | 14 | 13 | 15 | 15 | 19 | 14 |
| m7 | *An. stephensi* | 90 | 13 | 14 | 14 | 14 | 15 | 14 | 13 | 15 | 13 | 15 |
| m8 | *An. stephensi* | 95 | 18 | 18 | 16 | 14 | 14 | 16 | 15 | 16 | 14 | 14 |
| m9 | *An. stephensi* | 76 | 14 | 16 | 14 | 15 | 16 | 19 | 19 | 18 | 13 | 14 |
| m10 | *An. stephensi* | 80 | 14 | 13 | 14 | 14 | 15 | 14 | 16 | 16 | 14 | 14 |
| m11 | *An. stephensi* | 64 | 15 | 14 | 15 | 13 | 16 | 15 | 15 | 16 | 15 | 14 |
| m12 | *An. stephensi* | 81 | 13 | 16 | 16 | 14 | 15 | 15 | 14 | 13 | 13 | 16 |

Sporozoites were quantified on day 12 post feeding in salivary glands of mosquitoes that remained from the batch where high oocysts were detected during midgut dissection on day 7 post feeding and categorized into four (with a grade from 1-4) following protocol reported before (Joshi, Choochote et al. 2009). Table 7 depicts the detail results for each *An. arabiensis* and *An. stephensi* dissected (rows) with representative pictures in figure 1A-1B and summary in 1C.

**Table 7. Sporozoite quantification in paired membrane feeding experiments using colony *An. arabiensis* and wild caught *An. stephensi***

| *An. arabiensis* | | | | |
|---|---|---|---|---|
| **id** | **sporozoite** | | **id** | **sporozoite** |
| AD79-R01-1 | 2 | | AD86-R01-5 | 0 |
| AD79-R01-2 | 0 | | AD86-R01-6 | 0 |
| AD79-R01-3 | 0 | | AD86-R01-7 | 0 |
| AD79-R01-4 | 2 | | AD86-R01-8 | 4 |
| AD79-R01-5 | 0 | | AD86-R01-9 | 0 |
| AD79-R01-6 | 1 | | AD88-R01-1 | 2 |
| AD79-R01-7 | 4 | | AD88-R01-2 | 3 |
| AD79-R01-8 | 0 | | AD88-R01-3 | 4 |
| AD79-R01-9 | 0 | | AD88-R01-4 | 4 |
| AD79-R01-10 | 0 | | AD88-R01-5 | 4 |
| AD79-R01-11 | 0 | | AD88-R01-6 | 4 |
| AD79-R01-12 | 0 | | AD88-R01-7 | 0 |
| AD82-R01-1 | 0 | | AD88-R01-8 | 0 |
| AD82-R01-2 | 1 | | AD88-R01-9 | 1 |
| AD82-R01-3 | 0 | | AD88-R01-10 | 0 |
| AD82-R01-4 | 1 | | AD88-R01-11 | 4 |
| AD82-R01-5 | 0 | | AD88-R01-12 | 3 |
| AD82-R01-6 | 2 | | AD88-R01-13 | 4 |
| AD82-R01-7 | 1 | | AD88-R01-14 | 0 |
| AD82-R01-8 | 0 | | AD88-R01-15 | 4 |
| AD82-R01-9 | 0 | | AD88-R01-16 | 0 |
| AD82-R01-10 | 0 | | AD88-R01-17 | 4 |
| AD85-R01-1 | 4 | | AD88-R01-18 | 0 |
| AD85-R01-2 | 4 | | AD88-R01-19 | 4 |
| AD85-R01-3 | 2 | | AD88-R01-20 | 0 |
| AD85-R01-4 | 3 | | AD88-R01-21 | 3 |
| AD85-R01-5 | 2 | | AD88-R01-22 | 0 |
| AD85-R01-6 | 4 | | AD88-R01-23 | 2 |
| AD85-R01-7 | 2 | | AD88-R01-24 | 0 |
| AD85-R01-8 | 4 | | AD88-R01-25 | 0 |
| AD85-R01-9 | 2 | | AD88-R01-26 | 0 |
| AD85-R01-10 | 0 | | AD88-R01-27 | 0 |
| AD85-R01-11 | 2 | | AD88-R01-28 | 4 |
| AD85-R01-12 | 4 | | AD88-R01-29 | 4 |
| AD85-R01-13 | 4 | | AD88-R01-30 | 0 |
| AD85-R01-14 | 4 | | AD89-R01-1 | 0 |
| AD85-R01-15 | 4 | | AD89-R01-2 | 0 |
| AD85-R01-16 | 2 | | AD89-R01-3 | 0 |
| AD85-R01-17 | 1 | | AD89-R01-4 | 0 |
| AD85-R01-18 | 3 | | AD89-R01-5 | 0 |
| AD85-R01-19 | 2 | | AD89-R01-6 | 0 |
| AD86-R01-1 | 4 | | AD89-R01-7 | 0 |
| AD86-R01-2 | 2 | | AD89-R01-8 | 0 |
| AD86-R01-3 | 0 | | AD89-R01-9 | 0 |
| AD86-R01-4 | 3 | | AD89-R01-10 | 0 |

| *An. stephensi* | | | | | | | |
|---|---|---|---|---|---|---|---|
| **id** | **sporozoite** | | **id** | **sporozoite** | | **id** | **sporozoite** |
| AD79-R01-1 | 4 | | AD85-R01-20 | 4 | | AD88-R01-28 | 4 |
| AD79-R01-2 | 3 | | AD85-R01-21 | 2 | | AD88-R01-29 | 4 |
| AD79-R01-3 | 4 | | AD85-R01-22 | 3 | | AD88-R01-30 | 4 |
| AD79-R01-4 | 2 | | AD85-R01-23 | 4 | | AD89-R01-1 | 1 |
| AD79-R01-5 | 0 | | AD85-R01-24 | 2 | | AD89-R01-2 | 4 |
| AD79-R01-6 | 0 | | AD85-R01-25 | 3 | | AD89-R01-3 | 0 |
| AD79-R01-7 | 4 | | AD85-R01-26 | 3 | | AD89-R01-4 | 0 |
| AD79-R01-8 | 4 | | AD85-R01-27 | 4 | | AD89-R01-5 | 0 |
| AD79-R01-9 | 1 | | AD85-R01-28 | 4 | | AD89-R01-6 | 0 |
| AD79-R01-10 | 4 | | AD85-R01-29 | 2 | | AD89-R01-7 | 0 |
| AD79-R01-11 | 4 | | AD86-R01-1 | 1 | | AD89-R01-8 | 0 |
| AD79-R01-12 | 0 | | AD86-R01-2 | 0 | | AD89-R01-9 | 0 |
| AD82-R01-1 | 2 | | AD86-R01-3 | 2 | | AD89-R01-10 | 4 |
| AD82-R01-2 | 2 | | AD86-R01-4 | 0 | | AD89-R01-11 | 4 |
| AD82-R01-3 | 4 | | AD86-R01-5 | 0 | | AD89-R01-12 | 0 |
| AD82-R01-4 | 2 | | AD86-R01-6 | 0 | | AD89-R01-13 | 0 |
| AD82-R01-5 | 3 | | AD86-R01-7 | 2 | | AD89-R01-14 | 0 |
| AD82-R01-6 | 0 | | AD86-R01-8 | 0 | | AD89-R01-15 | 4 |
| AD82-R01-7 | 0 | | AD88-R01-1 | 4 | | AD89-R01-16 | 4 |
| AD82-R01-8 | 3 | | AD88-R01-2 | 4 | | AD89-R01-17 | 0 |
| AD82-R01-9 | 0 | | AD88-R01-3 | 4 | | AD89-R01-18 | 0 |
| AD82-R01-10 | 2 | | AD88-R01-4 | 4 | | AD89-R01-19 | 0 |
| AD82-R01-11 | 1 | | AD88-R01-5 | 4 | | | |
| AD82-R01-12 | 0 | | AD88-R01-6 | 4 | | | |
| AD82-R01-13 | 2 | | AD88-R01-7 | 3 | | | |
| AD82-R01-14 | 0 | | AD88-R01-8 | 1 | | | |
| AD85-R01-1 | 2 | | AD88-R01-9 | 1 | | | |
| AD85-R01-2 | 4 | | AD88-R01-10 | 0 | | | |
| AD85-R01-3 | 0 | | AD88-R01-11 | 3 | | | |
| AD85-R01-4 | 2 | | AD88-R01-12 | 1 | | | |
| AD85-R01-5 | 1 | | AD88-R01-13 | 4 | | | |
| AD85-R01-6 | 4 | | AD88-R01-14 | 4 | | | |
| AD85-R01-7 | 4 | | AD88-R01-15 | 4 | | | |
| AD85-R01-8 | 2 | | AD88-R01-16 | 4 | | | |
| AD85-R01-9 | 1 | | AD88-R01-17 | 4 | | | |
| AD85-R01-10 | 3 | | AD88-R01-18 | 4 | | | |
| AD85-R01-11 | 4 | | AD88-R01-19 | 1 | | | |
| AD85-R01-12 | 4 | | AD88-R01-20 | 1 | | | |
| AD85-R01-13 | 3 | | AD88-R01-21 | 4 | | | |
| AD85-R01-14 | 4 | | AD88-R01-22 | 4 | | | |
| AD85-R01-15 | 4 | | AD88-R01-23 | 4 | | | |
| AD85-R01-16 | 2 | | AD88-R01-24 | 3 | | | |
| AD85-R01-17 | 4 | | AD88-R01-25 | 4 | | | |
| AD85-R01-18 | 4 | | AD88-R01-26 | 4 | | | |
| AD85-R01-19 | 3 | | AD88-R01-27 | 4 | | | |

A

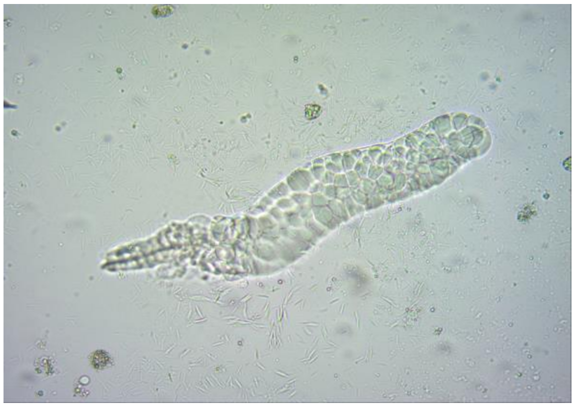

B

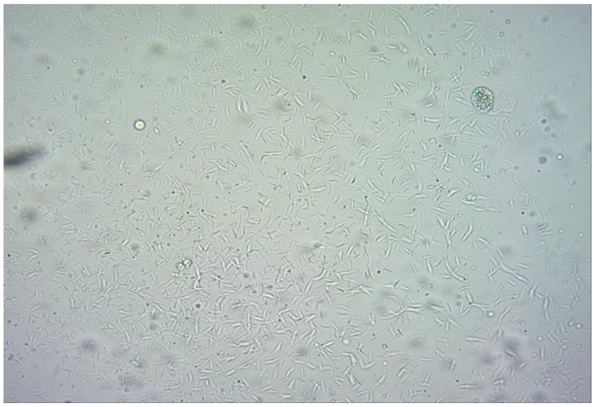

C

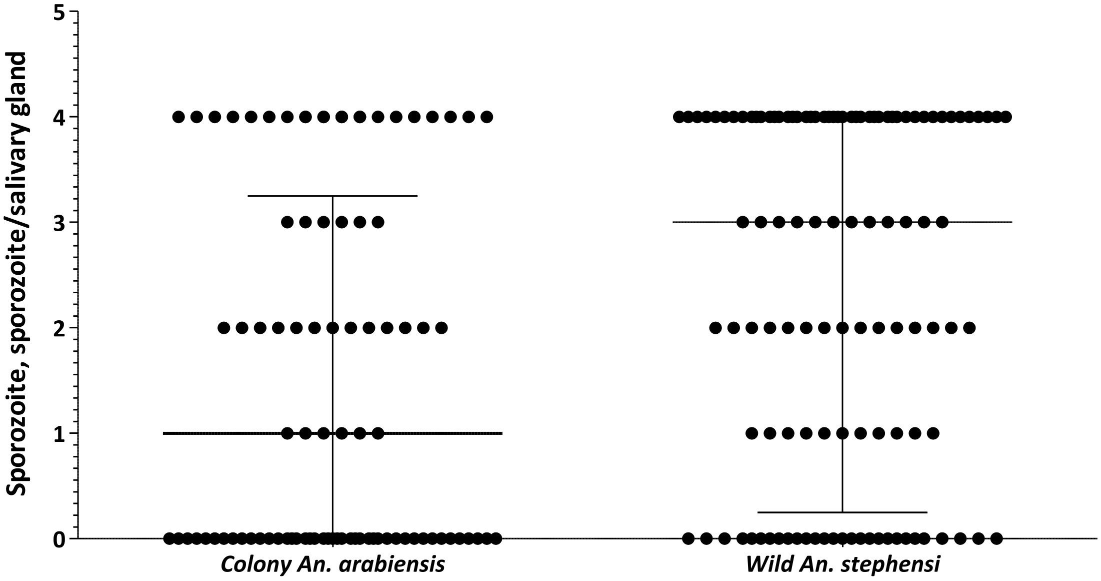


**Figure 1. Sporozoites detected from salivary glands of infected mosquitoes.** Dissection was done on day 12 post infection and picture was taken at 20X objective magnification. Salivary glands were graded as $1^+$ (when 1–10 sporozoites detected), $2^+$ (11–100 sporozoites), $3^+$ (101–1,000 sporozoites) and $4^+$ (above 1,000 sporozoites).

**Malaria incidence trend in the last five years in Awash town and its surroundings**

Weekly malaria surveillance data (total tested by microscopy/RDT and confirmed malaria by species) were obtained through the Ethiopian Public Health Institute at the Federal Ministry of Health for the years from 2015 to 2019. Test for trend of annual parasite incidence/proportion of infections across the five years defined by months was tested using the nonparametric test for trend across ordered groups (Cuzick 1985) after missing data points were imputed using mdesc command in STATA.

A total of 33,420 suspected cases were diagnosed for malaria in the past five years in Awash town of whom 3,328 (10.0%) were confirmed to have either *P. falciparum* (5.5%) or *P. vivax* (3.9%) infection. Overall, the API was higher in the surrounding of the town and has shown a non-significant decline over the previous years both in Awash town and its surrounding. In Awash town, there is an overall non-significant decline in total cases (z=-1.72; *P*=.086) which was mainly attributable to *P. vivax* (z=-3.99; *P*=.001) that aligns with the increase in the proportion of infections due to *P. falciparum* (z=3.15; *P*=.002) although there was no significant change in *P. falciparum* cases (z=-1.09; *P*=.277) (Figure 2). The proportion of *P. falciparum* infections has increased over the years in Awash town from 54.7% in 2015 to 76.4% in 2019 and remained stable in the surrounding.

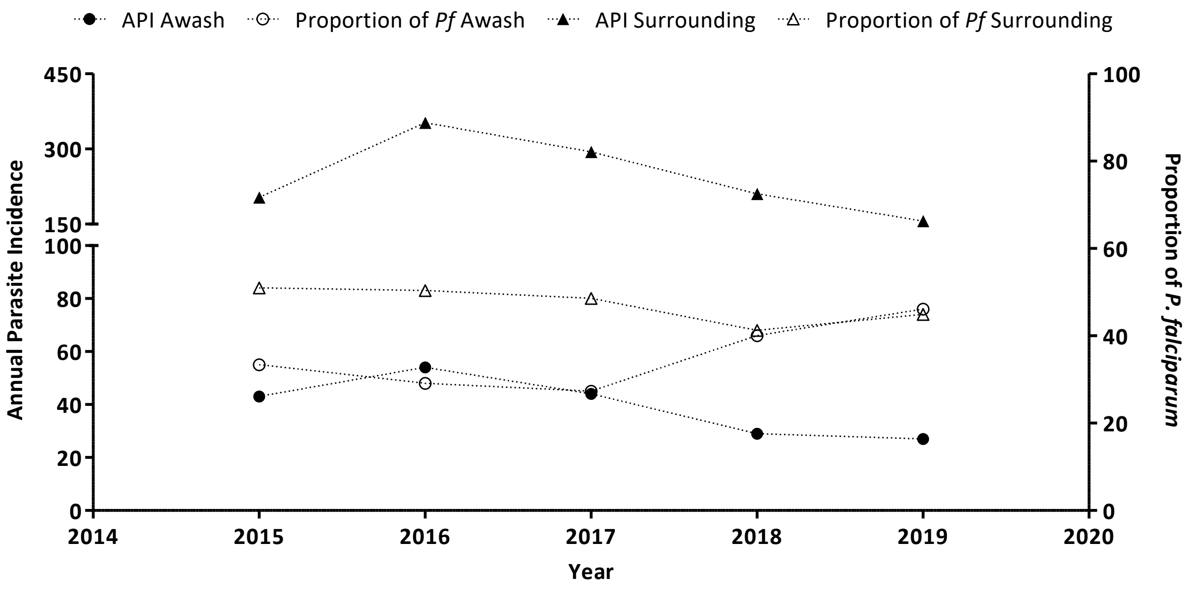


**Figure 2. Trend of annual parasite incidence and proportion of *P. falciparum* infections between January 2015 and September 2019.** Indicated on the left Y-axis is the annual parasite incidence per 1000 persons with the proportion of P. falciparum infections in the right Y-axis and years in the X-axis. In circles are the API (filled) and proportion (unfilled) of *P. falciparum* infections from Awash Sebat Kilo town. Filled triangles are API from Fentale district together with proportion of *P. falciparum* infections represented by unfilled triangles.
